# Diet-dependent effects of kombucha on the gut microbiome and its neuroactive potential: Associations with reduced anxiety and depressive-like behaviors in mice

**DOI:** 10.64898/2026.05.07.718715

**Authors:** Noor E Huma, Samuel Davison, Kylene Guse, Carrie Walls, Stephanie Rutschke, April Sackett, Gabriel Blanco, Junwei Zhang, Chi Chen, Juan Pablo Damián, Christopher Faulk, Andres Gomez

## Abstract

Fermented foods are increasingly recognized for their health-boosting potential, yet the mechanisms involved are not fully resolved. Here, we tested whether kombucha reshapes the gastrointestinal microbiome and whether these changes are associated with stress-related behaviors under contrasting dietary backgrounds. Male C57BL/6 mice were fed either a total Western diet (TWD) or a control diet (CTRL) supplemented with kombucha or water three times weekly for seven weeks. Depressive-like and anxiety-related behaviors were evaluated using the forced swimming (FST) and marble burying tests (MBT). Ileum, cecum, and colon microbiomes were profiled via 16S rRNA, ITS2, and shotgun metagenomics, while feces and whole brains were profiled by LC-MS metabolomics. Serum cytokines were measured by ELISA. Results highlight diet-dependent effects of Kombucha on behavioral, microbial and metabolic outcomes. Kombucha reduced immobility in the FST under both diets, whereas fewer marbles buried were observed only under TWD. Kombucha intake enriched Bifidobacterium pseudolongum in the ileum under CTRL and TWD diets, while cecal microbial functions related to amino acid metabolism were stimulated mainly under CTRL. Only CTRL mice receiving kombucha showed higher fecal acetate and butyrate together with lower fecal levels of neurochemically relevant amino acids, including glutamine, phenylalanine, tryptophan, and tyrosine. Under TWD, kombucha was associated with lower spleen weight and altered brain tryptophan/kynurenine profiles. These findings identify kombucha as a food intervention that can remodel gastrointestinal microbial and neuroactive metabolism in a diet depending manner. Associations with reduced depressive and anxiety-related behaviors are promising but warrant further exploration.

**Key Highlights:**

- Kombucha supplementation reshaped the mice gastrointestinal microbiome and its neuroactive potential
- Kombucha intake was associated reduced depressive and anxious like behaviors
- The potential of kombucha to modulate microbial, metabolic and behavioral outcomes may be dependent on subject dietary background

## INTRODUCTION

The gut microbiome plays a pivotal role in modulating the physiological landscape of humans and animals. One emerging area of investigation highlights the connections between dietary patterns, the gut environment and mental well being. This bidirectional communication between gut and brain is hypothesized to impact behavior, cognition, and emotional regulation (1,2) through several mechanisms. Some of the most widely studied include modulation of tryptophan (3) and serotonin metabolism (4,5) and production of γ-aminobutyric acid (GABA) (6) by the gut microbiome, effects of short-chain fatty acids (SCFAs) on blood brain barrier (BBB) integrity (7,8) and immune signaling pathways that shape neural plasticity and stress responses (9). Microbial regulation of gut-brain signaling has motivated a growing body of research on dietary strategies that target the gut environment to influence behavior and stress-related issues. For example, prebiotics and probiotics have been proposed as effective nutritional strategies to restore microbial balance and improve mental health (10–12). As mental health issues become a global public health problem (13), dietary and supplement-based approaches are increasingly recognized as widely accessible strategies for supporting psychological well-being.

Among the dietary interventions gaining increasing attention today, fermented foods have emerged as particularly promising, offering both physiological and psychological benefits through modulation of the gut microbiome and its downstream signaling pathways (14,15). Fermented foods such as kombucha, kefir, yogurt, kimchi, and sauerkraut contain probiotic microorganisms, organic acids, polyphenols, and bioactive metabolites that shape immune, metabolic, and vagal pathways connecting the gut environment and brain (16,17). Historically, these foods have been central to traditional healing practices aimed at restoring bodily and emotional balance (18,19), an idea now supported by advances in microbiome science and neurobiology (20).

Kombucha, a plant based fermented tea produced by a symbiotic culture of bacteria and yeast (SCOBY) and that originated around 2,200 years ago in China during the Qin Dynasty (21), has increasingly gained scientific interest for its purported benefits on health. Kombucha contains a symbiotic culture of live bacteria and fungi, plus a rich array of organic acids (mainly acetic and gluconic) and tea-polyphenols, which have the potential to beneficially influence immune and metabolic signaling (22,23). Such claims have raised Kombucha’s popularity and widespread consumption in western societies, in part due to a growing health and wellness market. Yet, studies that address kombucha’s potential to modulate the gut microbiome and its neuroactive functions remain scant. This knowledge gap is in contrast to the majority of existing reports demonstrating the neuroprotective potential of dairy fermented foods such as kefir and yogurt (17,24–27). Given substantial differences in substrate chemistry and microbial ecology found between dairy and plant ferments, kombucha may likely induce different pro/pre-biotic effects on the gut environment, including diverse metabolic and immune outcomes.

In this study, we hypothesized that kombucha consumption reshapes the gastrointestinal microbiome and its neuroactive metabolic potential, and that these changes are associated with modulation of stress-related behaviors. We further predicted that kombucha-associated effects on gut ecology, neurochemistry, and behavior would differ across dietary backgrounds. To test this, we exposed male C57BL/6 mice to either a standard control diet or a total Western diet to determine whether kombucha-associated responses are robust across distinct nutritional baselines or instead depend on a metabolically stressful dietary context. Because only male mice within a defined age range were included, the study was designed to evaluate diet-context effects under controlled biological conditions rather than to make sex or age-generalizable inferences. We profiled gut inter-kingdom ecology along the gastrointestinal tract using 16S rRNA, ITS2, and shotgun metagenomic sequencing, and characterized fecal and whole-brain metabolite profiles by LC-MS/MS. We also assessed spleen weight and serum cytokines as markers of systemic immune-related status. A schematic overview of the experimental design and sample collection workflow is shown in **Figure 1**.

**Figure 1.**
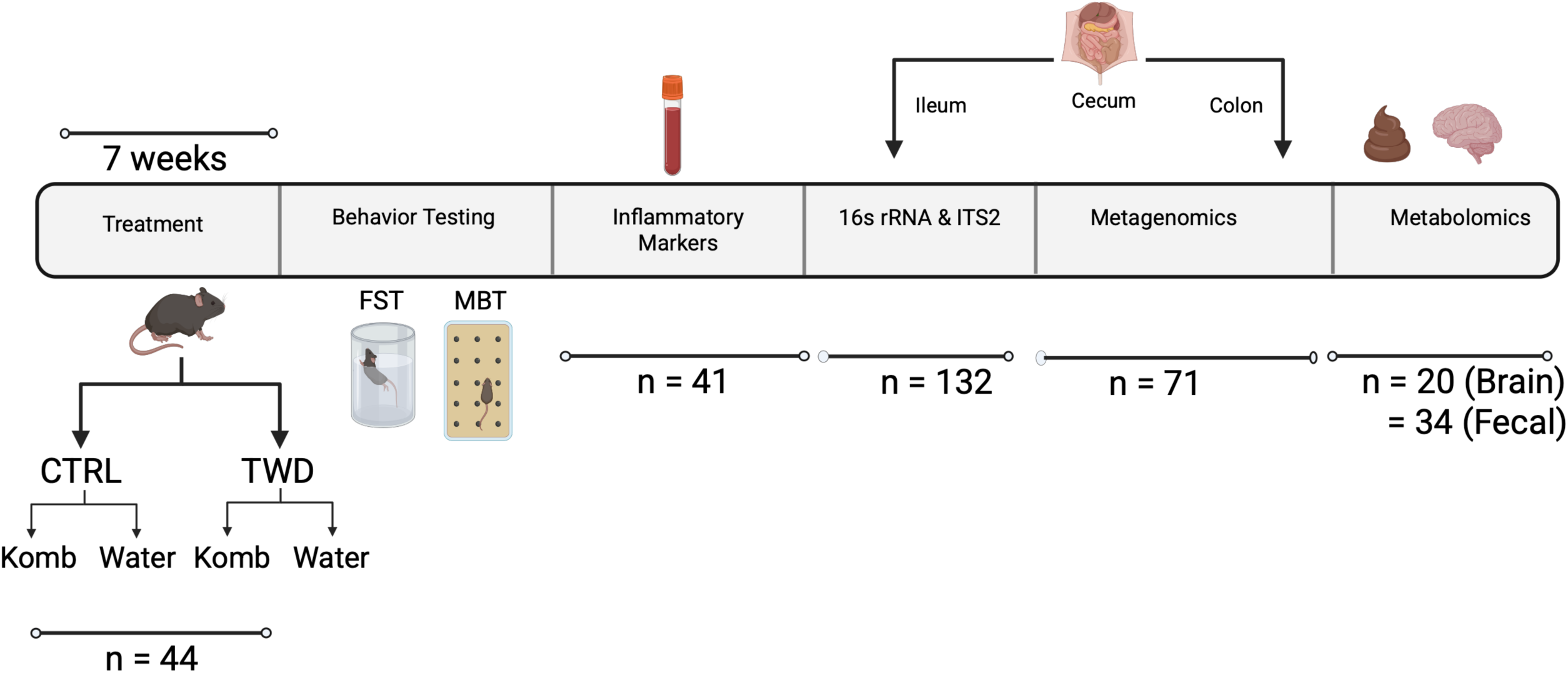
Experimental design and analytical workflow. Male C57BL/6 mice were fed either a control diet (CTRL) or a total Western diet (TWD) and supplemented with kombucha or water for 7 weeks. Endpoints included behavioral testing, serum inflammatory marker analysis, gastrointestinal 16S rRNA, ITS2, and shotgun metagenomic profiling, and fecal and whole-brain metabolomics. Created with BioRender.

## RESULTS

After a 7-week intervention in which male C57BL/6 mice were fed either a control diet (CTRL) or a total Western diet (TWD) and supplemented with kombucha (K) or water (W), kombucha was associated with behavioral, microbial, and metabolic changes across multiple host compartments. These included altered FST and MBT responses, selective shifts in spleen weight and inflammatory markers, remodeling of gastrointestinal bacterial and fungal communities, changes in microbial functional potential, and modification of fecal and, to a lesser extent, whole-brain metabolite profiles. Notably, these kombucha-associated effects were strongly dependent on dietary background.

### Kombucha intake was associated with altered immobility and marble-burying behaviors

Mice exposed to kombucha showed significantly reduced immobility in the forced swim test (FST) under both dietary backgrounds (P < 0.05 for both CTRL and TWD; **Figure 2A**). In contrast, reduced marble burying in the marble burying test (MBT) was observed only in mice receiving kombucha under the TWD (P = 0.05; **Figure 2B**). The FST and MBT were used as screening measures of depressive-like and anxiety-related or compulsive-like behaviors, respectively.

**Figure 2.**
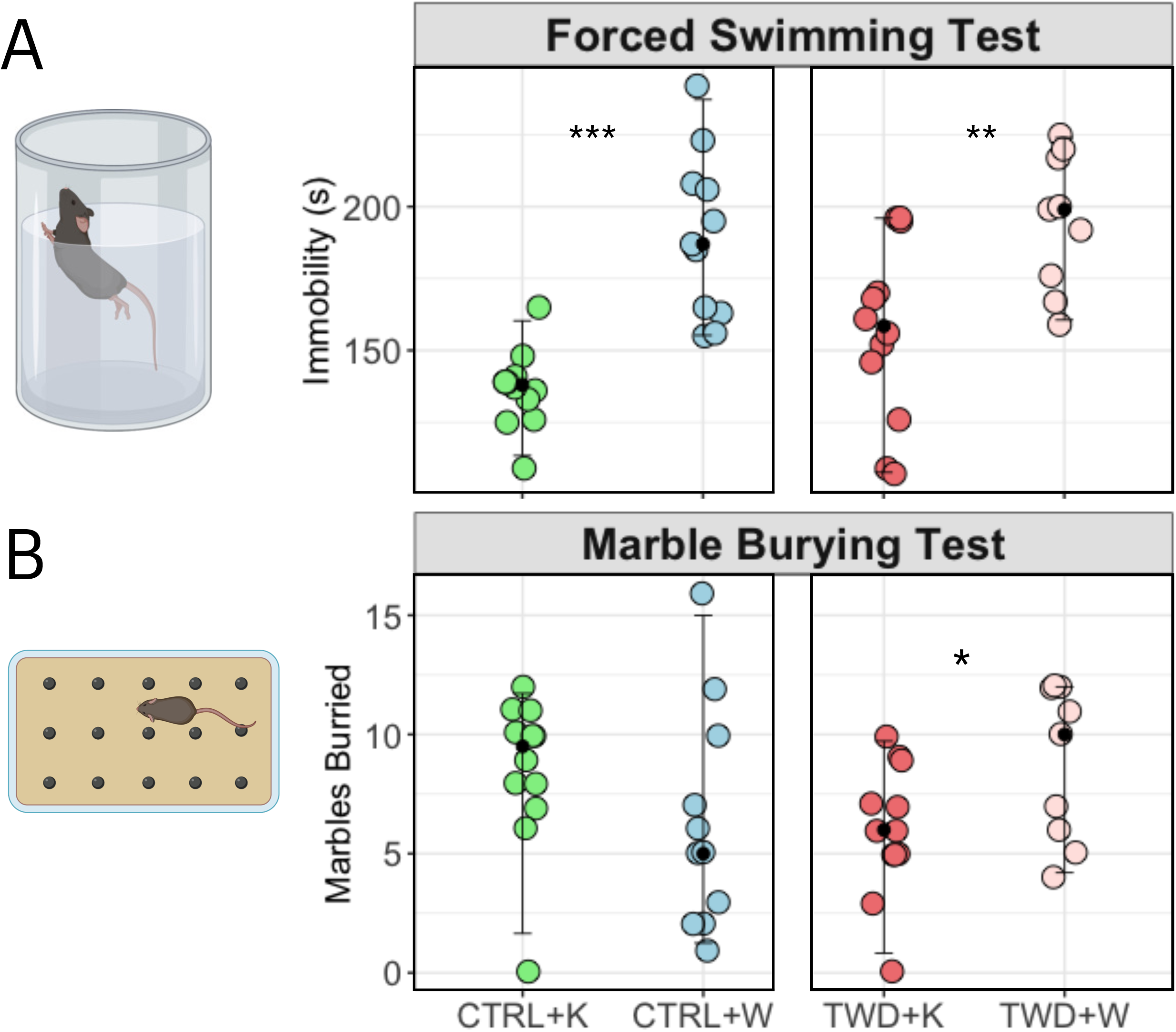
Kombucha was associated with altered depressive-like and marble-burying behaviors. (A) Forced swim test (FST) results, used here as a measure of depressive-like behavior. (B) Marble burying test (MBT) results, used here as a screening measure of anxiety-related and/or compulsive-like behavior. Significant differences were assessed within diet using Student’s *t*-test after testing for normality with the Shapiro–Wilk test. Data are shown as mean ± SEM. *P* < 0.05, P < 0.01, and *P* < 0.001. Behavioral icons were created with BioRender.

### Kombucha intake was associated with selective shifts in inflammation-related markers

Kombucha did not affect body weight under the TWD and showed only a trend toward increased body weight under the CTRL diet (P = 0.06, **Figure S1A**). Under the TWD, kombucha markedly reduced spleen weight (P = 0.0001), whereas under the CTRL diet spleen weight increased (P = 0.03, **Figure S1B**). Circulating GRO-α/CXCL1 was also elevated in CTRL-fed mice receiving kombucha (P = 0.03, **Figure S1C**), while IFNγ and IL-4 did not differ across groups.

### Kombucha intake induced diet- and site-dependent shifts in the gastrointestinal microbiome

We next examined kombucha-associated changes in bacterial and fungal communities along the ileum, cecum, and colon using 16S rRNA and ITS2 sequencing, respectively. Bacterial alpha diversity responses were site- and diet-dependent. For example, under TWD, kombucha reduced richness in the ileum and colon (Chao 1, P < 0.05), whereas under CTRL it tended to reduce richness in the cecum (P = 0.07) and significantly lowered Shannon diversity in the ileum (P = 0.0007; **Figure 3A-B**).

**Figure 3.**
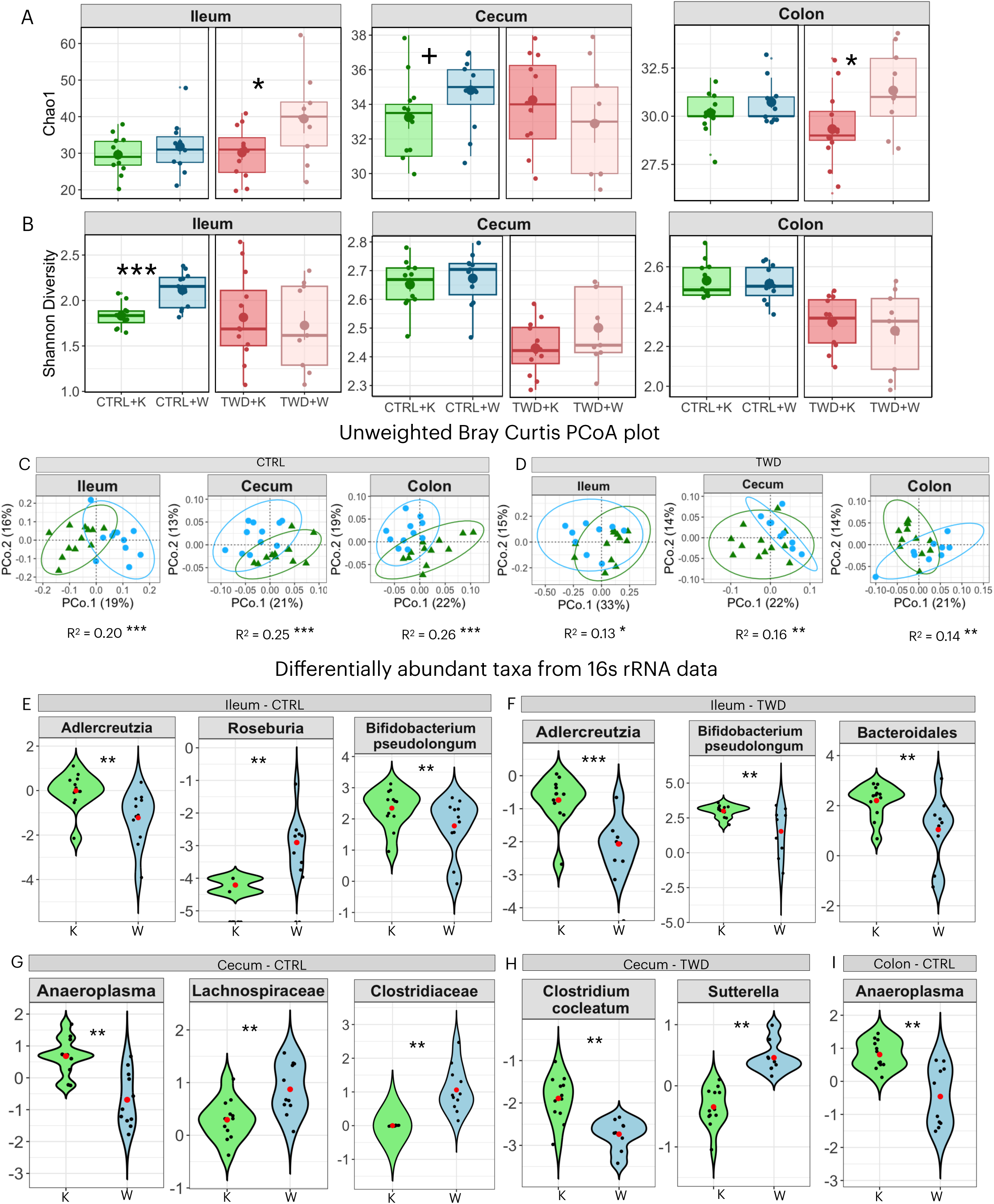
Kombucha induced diet- and site-dependent shifts in the gastrointestinal microbiome. (A-B) Alpha diversity of bacterial communities in the ileum, cecum, and colon, shown as Chao1 richness and Shannon diversity. Significance was assessed using the Mann–Whitney U test for within-diet comparisons. (C-D) Beta diversity of bacterial communities visualized by PCoA based on unweighted Bray-Curtis (Sørensen-equivalent presence/absence-transformed) distances; group differences were assessed by PERMANOVA. Ellipses represent 95% confidence intervals. (E-I) Differentially abundant bacterial taxa identified by MaAsLin3 (*P* < 0.05, *q* < 0.1). Significant differences are indicated as *P* < 0.05, P < 0.01, and *P* < 0.001; trends are indicated by + (*P* = 0.05-0.09).

Beta diversity analyses based on presence/absence-transformed Bray-Curtis distances (Sorensen) showed that diet was the dominant source of variation in bacterial community structure across all gut sites, with the strongest effects in the cecum and colon (P < 0.001; R² = 0.37-0.59, **Figure S2A**). Within-diet analyses, however, indicated that kombucha also contributed to bacterial community separation across all three sites, with stronger effects under CTRL (R² = 0.20-0.26, P < 0.001) than under TWD (R² = 0.13-0.16, P < 0.05; Figure 3C-D).

Weighted distances corroborated the dominant effect of diet, although kombucha-associated shifts within diet were less pronounced (**Figure S2B-D**). Intersection analysis indicated that most ASVs were shared across gut sites, but kombucha was associated with fewer unique and low-frequency intersection sets, particularly in the ileum under TWD, consistent with a more selective and less distinct site-specific bacterial community in this gut region (**Figure S3**).

At the taxonomic level, differential abundance analysis via MaAsLin3 showed that the most consistent bacterial response to kombucha was enrichment of *Bifidobacterium pseudolongum* and *Adlercreutzia* in the ileum under both diets (P < 0.01, q < 0.1; **Figure 3E-F**). Additional responses were diet- and site-specific, including increased *Anaeroplasma* in the cecum and colon under CTRL, enrichment of *Clostridium cocleatum* in the cecum under TWD, and reduced *Sutterella* in the cecum under TWD (**Figure 3G-I**). Other taxa, including *Staphylococcus* and *Akkermansia muciniphila*, showed nominal associations with kombucha consumption (P<0.05) but did not consistently remain significant after FDR correction at q<0.1 (**Figure S4**)

Fungal responses were more limited than bacterial ones. Kombucha increased fungal richness in the colon under CTRL but reduced diversity in the cecum under TWD (P<0.01, **Figure S5)**. Beta diversity analyses indicated that diet was the main driver of mycobiome structure, although effect sizes were substantially smaller than those observed for the bacteriome (P < 0.001; R² = 0.09-0.12, Figure **S6A-B**). Within-diet comparisons showed the clearest kombucha-related effects in the cecum and colon, again with smaller effect sizes than those observed for bacterial communities (P < 0.05; R² = 0.07-0.11, **Figures S5-S6**). Differential fungal signals were identified by indicator species analysis and Mann-Whitney U testing (Indicator value (IV) >0.5, P < 0.05), as MaAsLin3 with FDR correction did not detect any discriminating fungal taxa. Accordingly, associations between kombucha intake and higher abundance of fungal species such as *Saccharomyces cerevisiae* (colon) and *Vishniacozyma victoriae* (ileum) under CTRL should be considered preliminary and nominal (**Figure S7**).

### Kombucha intake altered the functional landscape of the gut microbiome

We performed shotgun metagenomic sequencing in a subset of samples (CTRL, n = 37; TWD, n = 34) to examine whether kombucha intake was associated with functional shifts across gastrointestinal sites (**Figure 4, Figure S8-S9 & supplementary file 1)**. The clearest FDR-supported functional responses were observed in the cecum under the CTRL diet, where kombucha was associated with enrichment of 19 microbial pathways (P<0.05, q<0.1, **Figure 4A**). These included L-lysine biosynthesis III, isoleucine biosynthesis III, valine biosynthesis and tetrapyrrole biosynthesis from glutamate, indicating altered microbial amino acid metabolic potential in this gut region. Because several of these pathways intersect with glutamate and branched-chain amino acid metabolism, the results are consistent with kombucha-associated shifts in microbial functions relevant to neuroactive precursor metabolism, although no direct functional validation was performed.

**Figure 4.**
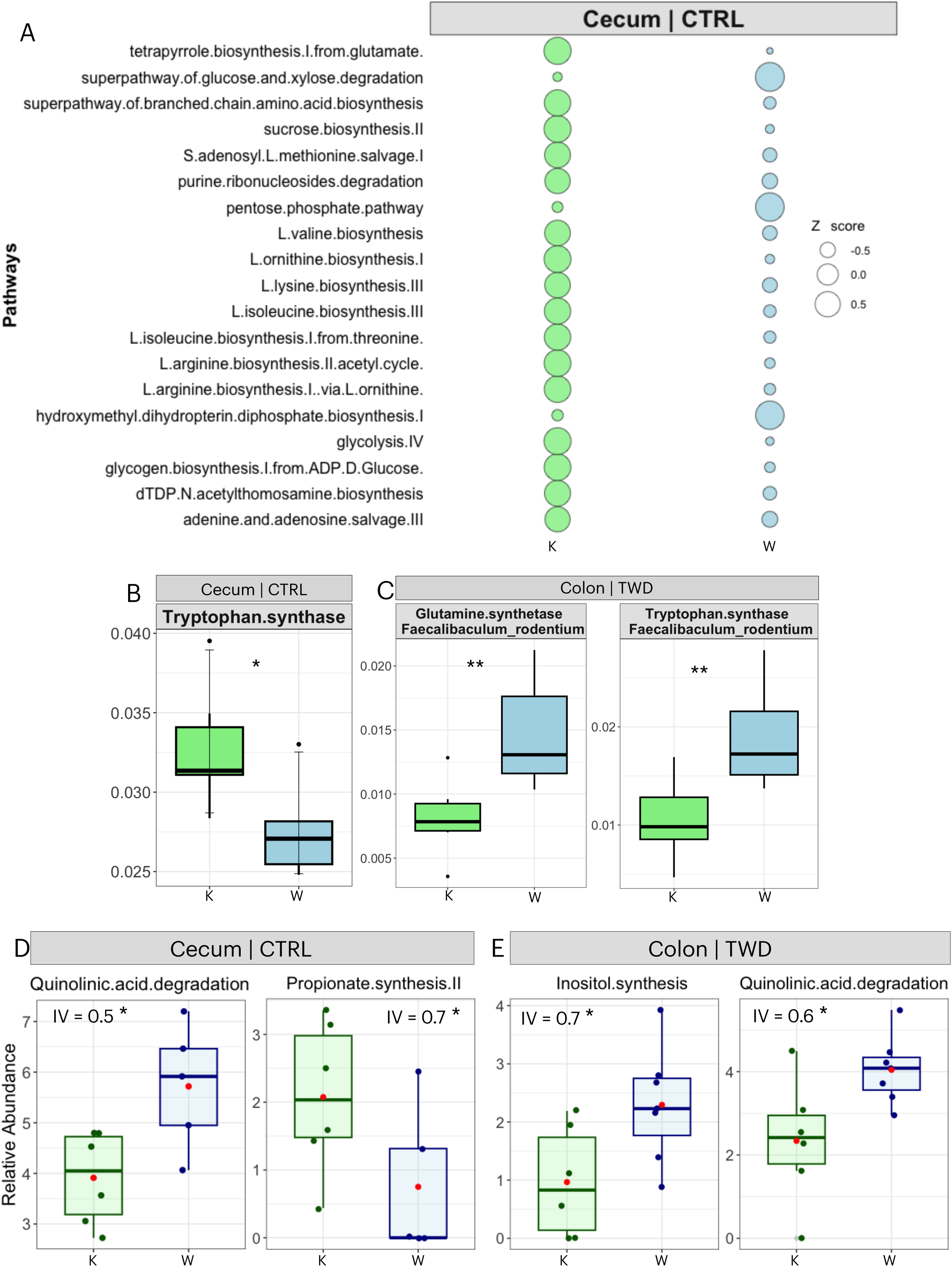
Kombucha altered the functional landscape of the gut microbiome. (A) Differentially abundant microbial pathways in the cecum under the CTRL diet identified by MaAsLin3 and visualized as a bubble plot (*P* < 0.05, *q* < 0.1). (B-C) Differentially abundant microbial genes in the cecum under CTRL and the colon under TWD identified by MaAsLin3 (*P* < 0.05, *q* < 0.1). (D-E) Differentially abundant gut-brain modules (GBMs) in the cecum under CTRL and the colon under TWD identified by indicator species analysis (IndVal; *P* < 0.05, indicator value > 0.6). Significant differences are indicated as *P* < 0.05, **P** < 0.01, and ***P*** < 0.001. Additional pathways and genes associated with kombucha intake are shown in Figures S8-S9 and Supplementary File 2.

Gene-level analyses also showed that the strongest FDR-supported signals occurred in the cecum under CTRL (**P < 0.05, q < 0.1, Figure 4B-C, Figure S9, and Supplementary File 1**). For example, kombucha was associated with higher abundance of tryptophan synthase in the cecum under CTRL, whereas glutamate synthase and tryptophan synthase assigned to *Faecalibaculum rodentium* were reduced in the colon under TWD (**Figure 4C**). Additional differentially abundant genes detected in the CTRL (cecum) and TWD (colon) included glutamate N-acetyltransferase and glutamate-5-semialdehyde dehydrogenase, further supporting site- and diet-specific effects on microbial amino acid metabolism. Although these amino acid-related pathways and genes are linked to the synthesis of neuroactive compounds, their enrichment should be interpreted as shifts in neuroactive potential rather than direct evidence of altered neurotransmission.

At the taxonomic level, no metagenomic species-level differences remained significant after FDR correction at q < 0.1. However, several nominal species-level associations (P < 0.05; q = 0.3-0.9) were directionally consistent with 16S-based genus-level results, including *Staphylococcus xylosus* and *S. nepalensis* in the ileum under CTRL and in the colon under both diets (**Supplementary File 2**). These species-level signals should therefore be considered exploratory.

To further examine microbiome functions with putative relevance to gut-brain signaling, we analyzed gut-brain modules (GBMs, **Figure 4D & 4E**). Differential GBM abundance was assessed using indicator species analysis (IV> 0.5) and corroborated by Mann-Whitney U testing as MaAsLin3 FDR correction did not detect any differentially abundant GBMs. Kombucha was associated with lower abundance of quinolinic acid degradation in the cecum under CTRL and in the colon under TWD, while propionate synthesis II was enriched in the cecum under CTRL (P < 0.05, **Figure 4D-E**). In combination with the gene-level findings above, these patterns are consistent with altered microbial functions related to tryptophan and short-chain fatty acid metabolism, but they do not establish directionality in vivo. Likewise, although propionate-related functions have been linked to blood-brain barrier biology (8), the present data do not indicate whether this enrichment translated into higher systemic propionate levels.

### Kombucha modulated fecal and whole brain metabolites

To further characterize metabolic features associated with kombucha intake, we performed targeted LC-MS metabolomics on fecal samples, including amino acids, SCFAs, and lactic acid, and on whole-brain samples for selected neuroactive metabolites (**Figure 5, S10-S12**). Fecal metabolite profiles were primarily diet-driven (PERMANOVA P=0.02, R²=0.99, **Figure S10A**), with kombucha-associated shifts evident only under the CTRL diet (P=0.01, R²=0.2, **Figure S10B**). CTRL-fed mice receiving kombucha also showed lower fecal metabolite richness than water controls (P = 0.03, **Figure S10D**).

**Figure 5.**
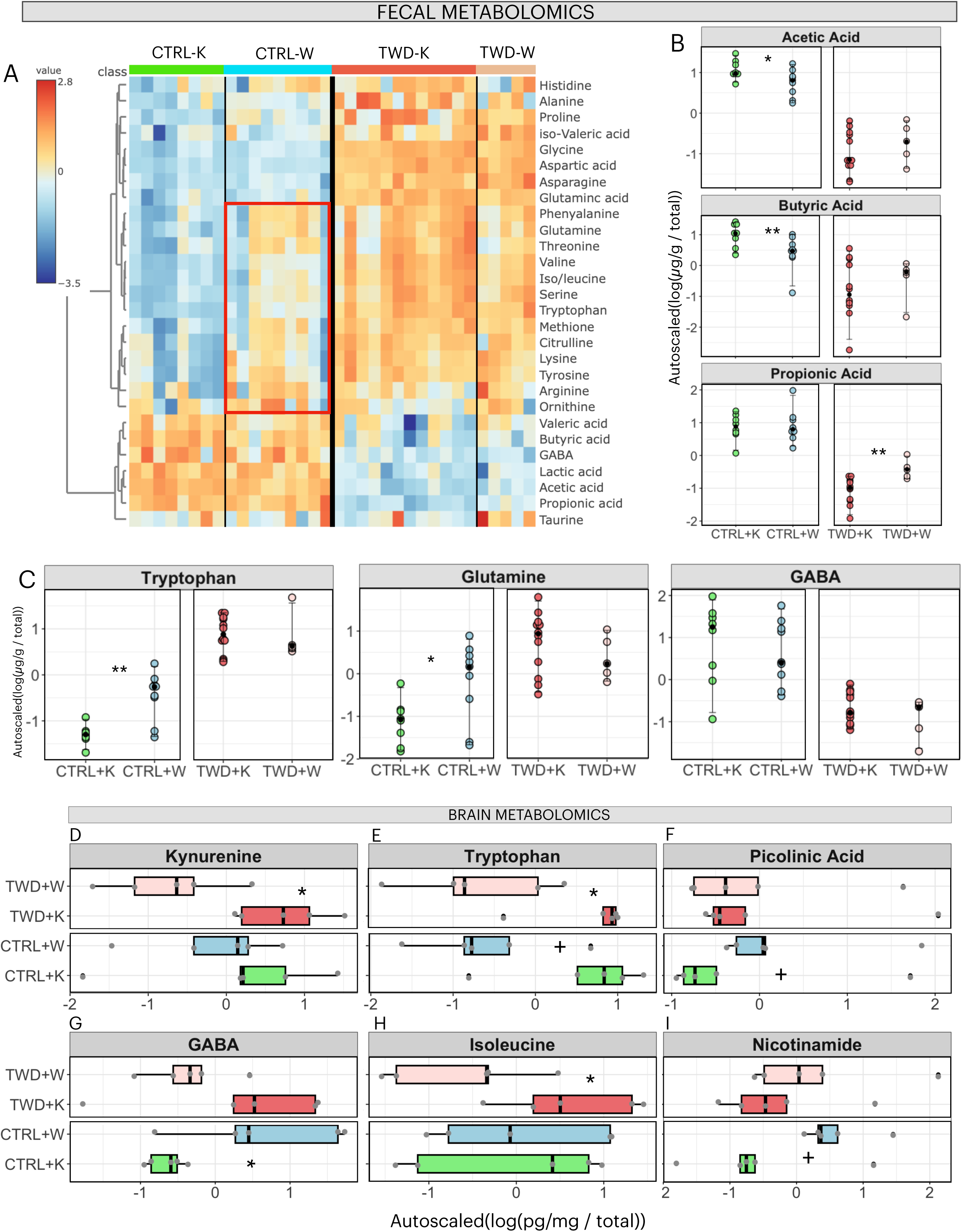
Kombucha was associated with distinct fecal and whole-brain metabolite profiles. (A) Heatmap showing relative abundance of fecal amino acids, lactic acid, and short-chain fatty acids (SCFAs) in mice fed CTRL or TWD and supplemented with kombucha or water. Heatmap clustering was based on normalized data, Euclidean distance, and Ward’s linkage using average metabolite abundance. (B-C) Beeswarm plots showing selected fecal SCFAs and amino acids. (D-I) Boxplots showing selected whole-brain metabolites. Significance for within-diet comparisons was assessed using the Mann–Whitney U test. *P* < 0.05, **P** < 0.01, and ***P*** < 0.001 indicate significant differences; + indicates a trend (*P* = 0.05-0.09). Metabolomics data were normalized by sum, log transformed, and autoscaled.

The fecal metabolome differed markedly by dietary background. TWD-fed mice showed higher abundance of most amino acids, whereas CTRL-fed mice were enriched in acetate, butyrate, propionate, lactic acid, GABA, and valeric acid (**Figure 5A-C**). Within CTRL, kombucha was associated with lower fecal abundance of several amino acids relevant to neuroactive metabolism, including glutamine, arginine, tryptophan, tyrosine, phenylalanine, threonine, and branched-chain amino acids (q<0.1, P<0.05, **Figures 5A & 5C**). While all SCFAs were significantly depleted under TWD regardless of kombucha intake, kombucha was associated with higher fecal SCFAs, specifically acetate and butyrate under CTRL (P<0.05, **Figure 5B**). Together, these findings indicate diet-dependent shifts in fecal fermentation products and amino acid pools associated with kombucha intake. Abundance of other fecal metabolites can be found in **Figure S11.**

Targeted analysis of whole-brain metabolites showed no significant multivariate effect of diet or kombucha when all measured metabolites were considered together (**Figure S12A-D**). At the individual metabolite level, however, kombucha was associated with higher kynurenine and tryptophan under TWD (P < 0.05, **Figure 5D-E**), with a borderline increase in tryptophan under CTRL (P = 0.09). In contrast, whole-brain GABA was lower under CTRL with kombucha exposure (P = 0.05). Borderline differences were also observed for isoleucine under TWD and nicotinamide under CTRL (P = 0.09; **Figure 5H-I**), while other measured brain metabolites did not differ significantly (**Figure S12E-F**). These results support diet-dependent associations between kombucha intake and host metabolite profiles, while indicating that fecal and whole-brain signatures do not shift in parallel across all metabolite classes.

### Exploratory multi-omic integration revealed diet-dependent association patterns under kombucha exposure

To obtain an integrative view of microbiome-metabolite relationships associated with kombucha intake, we performed an exploratory multi-omic correlation analysis using metagenomic features, including taxa, genes, and gut-brain modules (GBMs), together with fecal metabolites. Compositionally corrected positive Spearman correlations were retained at P < 0.05, Spearman’s rho > 0.6; these results should therefore be interpreted as exploratory and hypothesis-generating rather than confirmatory. Distinct diet-dependent association patterns were observed in mice receiving kombucha under CTRL versus TWD (**Figure 6A-B**).

**Figure 6.**
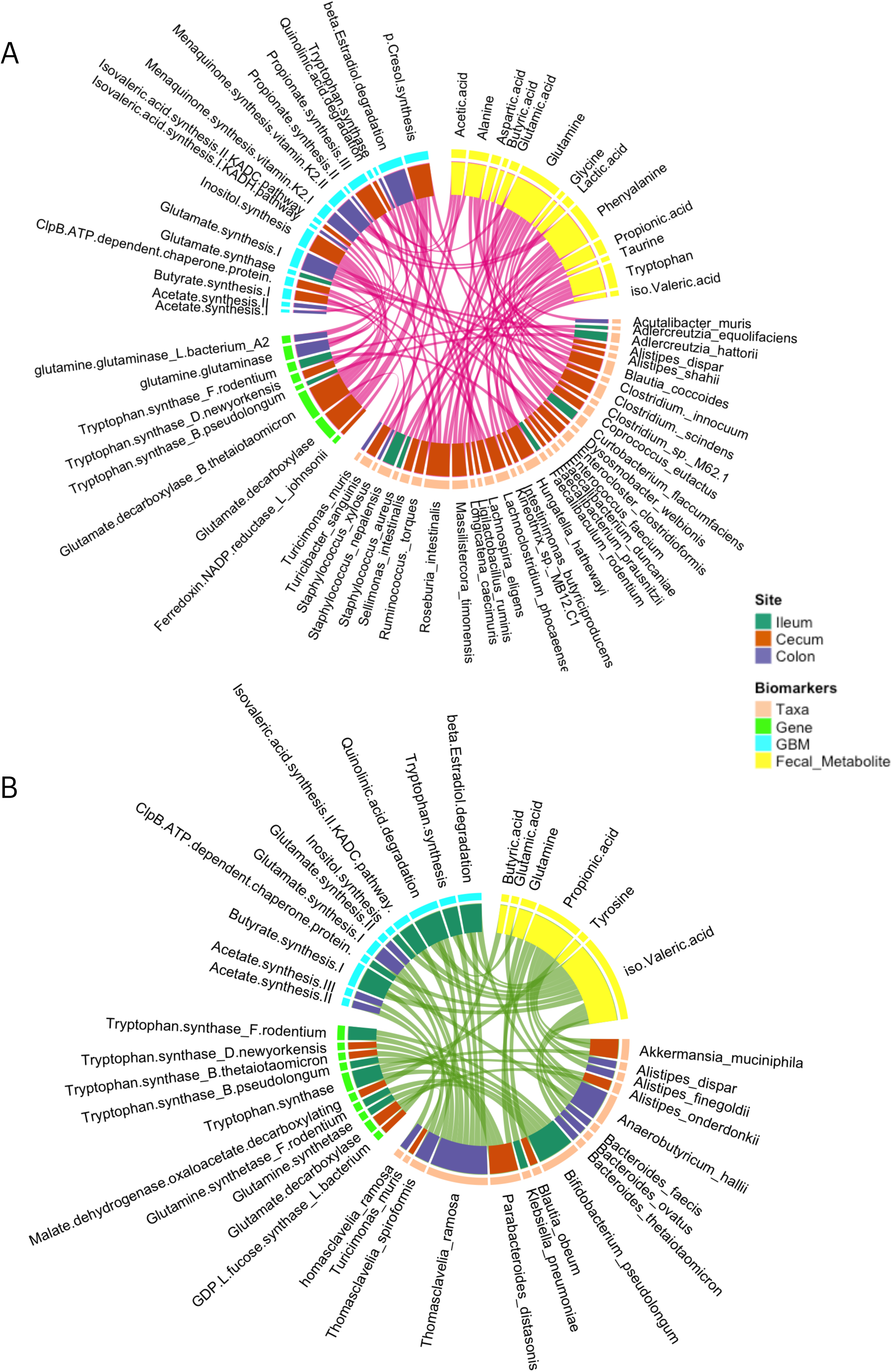
Exploratory microbiome-metabolite associations under CTRL and TWD. (A-B) Chord diagrams showing exploratory associations between metagenomic features (taxa, genes, pathways, and gut-brain modules) and fecal metabolites in mice receiving kombucha under CTRL (A) or TWD (B). Only positive associations are shown (P < 0.05, Spearman’s rho > 0.6). Because these multi-omic associations were exploratory, they should be interpreted as hypothesis-generating.

Under the CTRL diet, the correlation structure was more extensive and was concentrated primarily in the cecum. Several associations involved taxa commonly linked to short-chain fatty acid metabolism, including *Turicibacter sanguinis* with acetate and propionate, and *Clostridium sp*. with propionate. Additional associations included phenylalanine with *Staphylococcus aureus* in the ileum and tryptophan with *Ruminococcus torques* in the cecum. While these links are biologically relevant in the context of intestinal amino acid metabolism, they should not be interpreted as evidence that these taxa directly generate the corresponding luminal amino acid pools in this dataset. All three major SCFAs also showed positive associations with the glutamine-glutaminase gene in the colon, further supporting coordinated shifts between microbial functional potential and fecal amino acid profiles under CTRL.

Under the TWD, fewer overall associations were detected, but several linked taxa with SCFA-related functions and GBMs. These included associations of *Anaerobutyricum hallii* with the butyrate synthesis I GBM in the cecum and colon, *Alistipes dispar* and *A. finegoldii* with propionate in the colon, and *Akkermansia muciniphila* with propionate in the cecum. We also observed associations between *Thomasclavelia ramosa* and GBMs linked to acetate, butyrate, glutamate, and tryptophan metabolism. Compared with CTRL, these TWD associations appeared more concentrated in the ileum, consistent with the broader site-specific patterns described above, but they remain correlational and do not establish functional directionality in vivo. Additional correlating biomarkers from metagenomic features and fecal metabolome can be seen in **supplementary file 3**.

A separate exploratory analysis identified positive correlations between ileal *Akkermansia muciniphila* and whole-brain GABA (P = 0.02) and isoleucine (P = 0.01) under TWD (**Figure S13**). These observations are notable given reports on the beneficial roles of *A. muciniphila* in intestinal function, but these results cannot establish that intestinal abundance of this taxon influences brain metabolite levels directly.

To further characterize community-level organization, we examined co-abundance networks across gut sites under both diets (**Figure S14**). Network structure varied markedly by site and diet. The most visually distinct pattern was observed in the ileum under TWD, where kombucha exposure was associated with higher neighborhood connectivity and degree and lower average shortest path length relative to water controls. In contrast, under CTRL, kombucha was associated with opposite network traits, that is, lower ileal connectivity and degree but higher path length. In the cecum, kombucha increased neighborhood connectivity while reducing degree and path length under both diets, whereas in the colon kombucha intake was generally associated with lower connectivity and degree and higher path length. Together, these network results indicate that kombucha is associated with site and diet-dependent restructuring of multi-omic co-abundance patterns, although the biological consequences of these topological shifts remain unresolved.

## DISCUSSION

Together, our findings indicate that kombucha was associated with site- and diet-dependent shifts in the gut microbiome at both compositional and functional levels. These shifts included enrichment of microbial functions related to short-chain fatty acid (SCFA) metabolism and amino acid pathways relevant to neuroactive precursor metabolism, alongside reduced depressive-like and anxiety-related behaviors and selective changes in inflammation-related markers. An important aspect of the study design was the inclusion of both CTRL and TWD backgrounds, which revealed that kombucha-associated responses were not uniform but strongly conditioned by dietary context. A schematic summary of these findings and potential mechanistic implications for further testing is shown in **Figure S15**.

Kombucha is rich in acetic acid, gluconic acid, and tea polyphenols (28), which may act as ecological filters that favor niche-adapted bacteria and fungi along the gastrointestinal tract. In this context, kombucha did not broaden microbial diversity uniformly, but instead intake was associated with reduced diversity in a site- and diet-dependent manner. These results are broadly consistent with reports that fermented food intake can promote selective microbial restructuring rather than generalized diversification (29–31). Our data extend this concept by suggesting that such selectivity differs across gastrointestinal regions, including the ileum, a gut site with prominent immune activity.

One of the most consistent taxonomic responses to kombucha was enrichment of Bifidobacterium pseudolongum in the ileum under both CTRL and TWD. Previous studies have shown that B. pseudolongum can enhance IL-22 signaling and strengthen intestinal barrier function through the PPARγ/STAT3 pathway (32,33). Improved epithelial integrity could, in principle, influence enterochromaffin cell function and host tryptophan handling (34). B. pseudolongum has also been linked to regulation of systemic tryptophan metabolism (35). In our dataset, B. pseudolongum co-occurred with genes related to tryptophan and glutamine/glutamate metabolism, a pattern consistent with a possible role in modulating microbial functions relevant to neuroactive potential. However, these associations cannot be interpreted as direct evidence that this taxon drives systemic serotonin or GABA pools in vivo.

The enrichment of pathways linked to L-lysine biosynthesis and tetrapyrrole biosynthesis from glutamate in the cecum under kombucha exposure is also consistent with altered microbial amino acid metabolism in this gut region (36,37). Coupled with lower fecal glutamine under CTRL, these findings may reflect redistribution or altered microbial utilization of amino acid pools relevant to neuroactive metabolism (38). At the same time, the colon showed a different functional profile, with fewer glutamate-related signals, further underscoring that kombucha-associated responses differed across gut sites.

A more difficult result to interpret was the observation that, under the CTRL diet, kombucha was associated with lower whole-brain GABA while fecal GABA remained unchanged. This dissociation highlights the challenge of linking peripheral and central GABA pools directly. Gut-derived GABA can influence host physiology through local signaling and potentially through vagal pathways (39), and both lower systemic and cortical GABA have been associated with depressive disorders (40–42). However, our results do not explain why CTRL-fed mice showed lower whole-brain GABA, nor why TWD mice showed an opposite, albeit non-significant, trend. Moreover, GABA receptor expression and GABAergic tone vary across brain regions (39,43), so whole-brain metabolomics may mask region-specific changes. Thus, the present data should only be interpreted as evidence of compartment-specific dissociation rather than a straightforward kombucha-induced increase in GABAergic signaling.

Additional evidence that kombucha was associated with microbial functions relevant to host neurochemistry came from gut-brain module analyses, where lower abundance of quinolinic acid degradation was detected in the cecum and colon under kombucha exposure. This pattern, together with enrichment of tryptophan synthase in selected gut regions, is consistent with altered microbial functions related to tryptophan metabolism (44). A psychobiotic diet enriched in fiber, prebiotics, and fermented foods has previously been associated with lower quinolinic acid in humans and reduced perceived stress (45). Even so, our data do not establish pathway directionality or demonstrate that kombucha shifts tryptophan metabolism away from the kynurenine pathway toward serotonin production. This is particularly important given that neither TPH1 nor GAD was differentially abundant, and systemic inflammatory markers such as IFN-γ were not significantly altered (46). It is also important to note that differential abundance of GBMs should be tested in a better powered dataset.

Our fecal and brain metabolomics results further support the idea that kombucha is associated with diet-dependent reorganization of tryptophan-related metabolite pools. Under CTRL, kombucha was associated with lower fecal tryptophan and a borderline increase in whole-brain tryptophan. This pattern may reflect altered partitioning of tryptophan between the gut lumen and host tissues, potentially through changes in microbial utilization, host absorption, or both (47). Because serotonin and kynurenine themselves were not comprehensively resolved across all compartments, the results should be interpreted as evidence of altered tryptophan handling rather than direct proof of altered serotonergic signaling, which would require systemic and mechanistic validation.

Several microbial changes also raise the possibility that kombucha may influence host barrier-related functions. Under CTRL, kombucha was associated with higher abundance of Saccharomyces cerevisiae in the colon; this fungus has previously been linked to increased expression of occludin and claudin-1 in mice (48). Kombucha was also associated with increased Anaeroplasma in the cecum and colon, and this genus has been implicated in TGF-β-related immunoregulatory pathways (49). In our study, however, fungal associations were nominal and did not survive FDR-based differential abundance testing, so these observations should be viewed as preliminary. Likewise, the barrier-related relevance of *Anaeroplasma* and *S. cerevisiae* remains inferential here because tight junction proteins and mucosal immune markers were not directly measured.

A similar degree of caution is warranted for *Staphylococcus*. Although *Staphylococcus* increased across several gut regions under kombucha exposure, the relevant species-level signals were nominal rather than FDR-supported. Some *staphylococci*, especially *Staphylococcus xylosus*, encode the aromatic amino acid decarboxylase sadA, which can convert aromatic amino acids into bioactive amines such as serotonin and dopamine precursors (47,50). However, because the species-level signals in our study were exploratory and because direct metabolic flux was not measured, these observations should be treated as hypothesis-generating. The same applies to the co-occurrence of *Clostridiaceae*-related signals and tryptophan synthase in the cecum, which is biologically suggestive but not mechanistically demonstrative (51).

SCFA-related findings were among the more consistent metabolic signals associated with kombucha, especially under CTRL. Kombucha exposure was associated with higher fecal acetate and butyrate under CTRL, while all SCFAs and lactic acid were significantly depleted under TWD regardless of kombucha intake. These findings point to a strong dietary-context effect on fermentation-associated responses. Butyrate is well known to influence mucosal integrity, immune signaling, and host gene regulation, including pathways relevant to gut-brain communication (47,52,53). Nonetheless, our data do not establish whether the observed fecal changes reflect increased production, altered absorption, or altered host utilization. Therefore, SCFA results are best interpreted as evidence of altered fermentation-related metabolic output rather than proof of downstream neuroprotective effects.

One of the central themes emerging from this study is that kombucha-associated effects differed substantially by both dietary background and gut site. Under CTRL, kombucha was more strongly associated with cecal and colonic functional-metabolic integration, whereas under TWD, the most distinctive patterns emerged in the ileum. Exploratory chord diagrams and co-abundance networks were consistent with this interpretation, but because these analyses relied on relatively permissive q-value thresholds and correlation structure, they should be considered hypothesis-generating rather than confirmatory. Even so, the observation that kombucha under TWD was associated with denser ileal multi-omic networks, together with reduced spleen weight and enrichment of taxa such as *B. pseudolongum* and *Akkermansia muciniphila*, supports the broader interpretation that dietary background shapes how fermented food intake is associated with host-microbiome organization.

The positive correlation between ileal *A. muciniphila* and brain GABA under TWD is also intriguing in this context. *A. muciniphila* encodes the glutamate decarboxylase system and can produce GABA under defined conditions (54), and it has also been implicated in host metabolic regulation, including isoleucine-related pathways (55). However, the association observed here does not demonstrate that ileal *A. muciniphila* directly influenced brain GABA or isoleucine levels. Rather, it highlights a potentially informative link between intestinal taxa and host metabolic signatures that warrants targeted mechanistic testing.

Behaviorally, kombucha reduced immobility in the forced swim test under both diets, whereas reduced marble burying was observed only under TWD. These results are consistent with antidepressant-like and anxiety-related behavioral shifts, but they should be interpreted within the limitations of the behavioral battery used (56–60). Importantly, the metabolic correlates of these behavioral changes differed by dietary context. Under CTRL, kombucha was associated with higher fecal acetate and butyrate and lower fecal amino acid pools, whereas under TWD it was associated with altered tryptophan/kynurenine patterns and lower spleen weight. Thus, rather than acting through a single conserved mechanism, kombucha may be associated with distinct microbiome-metabolite-behavior signatures and mechanisms depending on dietary baseline.

### Limitations of the study

Several limitations should be considered when interpreting these findings. First, kombucha is a complex fermented food matrix, and this study did not seek to isolate the specific microbial or chemical constituents responsible for the observed associations. Second, the sample size was modest and the intervention duration was relatively short. Third, behavioral inference relied primarily on the forced swim and marble burying tests; additional assays, including locomotor control measures, would strengthen interpretation. Fourth, only male mice were studied, limiting inference across sex. Fifth, whole-brain metabolomics cannot resolve region-specific neurochemical changes. Sixth, kombucha was administered by gavage, which improves dose control but does not fully model voluntary consumption and bypasses potential cephalic-phase responses. Seventh, several multi-omic integration results, including species-level metagenomic findings, fungal associations, and network/correlation patterns, were exploratory. Finally, causality cannot be inferred because no fecal microbiota transplantation, metabolite supplementation, depletion, or genetic perturbation experiments were performed. Future work integrating region-specific brain analyses, targeted barrier and immune markers, and direct causal tests will be required to define how kombucha-associated microbial and metabolic changes relate to host behavior.

### Conclusion

Kombucha intake was associated with reduced depressive-like behavior in mice and with diet-dependent remodeling of gastrointestinal microbiomes, microbial functions, and metabolite profiles. These shifts included changes in SCFA-related metabolism, amino acid pathways and metabolites relevant to neuroactivity (tryptophan and glutamate), and exploratory multi-omic signatures potentially relevant to gut-brain communication (summarized in **Figure S15**). Overall, the findings support kombucha as a fermented food matrix capable of reshaping host-associated gut ecology in a diet-sensitive manner, while motivating follow-up mechanistic studies to determine how such responses may be leveraged in precision nutrition and broader food-based strategies for mental well-being. If confirmed mechanistically, these data further suggest that probiotic or fermented drink interventions may be more effective when aligned with an individual’s dietary practices and baseline gut ecology rather than applied as one-size-fits-all therapies.

## MATERIALS AND METHODS

### Animals and housing

Male C57BL/6 mice were obtained post-weaning from a commercial vendor and acclimated for 1 week before the intervention. Mice were group-housed (4 mice/cage; 11 cages total) under a 12 h light/dark cycle (lights on at 08:00) with controlled temperature and humidity and ad libitum access to water. Animals were monitored daily for general health, and body weight was recorded weekly. All procedures were approved by the University of Minnesota Institutional Animal Care and Use Committee (Protocol #2105-39093A).

### Diets, experimental design, and kombucha administration

Mice were assigned to one of two dietary backgrounds: a control chow diet (CTRL; Teklad Global 2018) or a total Western diet (TWD; TD.110424), the latter selected to model a high-fat, high-carbohydrate dietary background with lower micronutrient density. Inclusion of both diets was intended to test whether kombucha-associated responses were consistent across distinct nutritional baselines and whether they differed under a metabolically stressful dietary context.

Within each diet, mice received either kombucha or water for 7 weeks, generating four groups: CTRL+K, CTRL+W, TWD+K, and TWD+W (n = 11/group). Kombucha was administered by oral gavage using a 24–22 gauge feeding needle. During the first 3 weeks, mice were dosed once daily for 5 days/week; thereafter, dosing frequency was reduced to 3 times/week until study completion. Stool was collected weekly throughout the intervention. Behavioral testing was performed at week 7, and endpoint tissue collection occurred 24 h after the final administration. Detailed kombucha preparation is provided in Supplementary File 4.

### Behavioral testing

Behavioral assays were performed from least to most stressful with at least 24 h recovery between tests. The forced swim test (FST) was used as a measure of depressive-like behavior and was conducted as previously described (61). Mice were placed individually in 13 cm diameter cylinders containing water at 22°C for 6 min, and immobility during the final 4 min was quantified using ANY-maze and verified by hand scoring.

The marble burying test (MBT) was used as a screening measure of anxiety-related and/or compulsive-like behavior (58–60,62). Mice were placed in a clear cage containing 5-10 cm of lightly tamped bedding with evenly spaced marbles, and the number of marbles buried to at least two-thirds of their depth was recorded at the end of the test. Because MBT alone does not provide a comprehensive assessment of anxiety, the results were interpreted conservatively as a screening behavioral readout.

### Specimen collection

Stool samples were collected weekly from weeks 1–7 and stored at −80°C. For metabolomics, fecal samples from weeks 5–7 were pooled within the mouse. At endpoint, approximately 50 µL of blood was collected from the saphenous vein into serum separator tubes, centrifuged at 1,200 × g for 10 min, aliquoted, and stored at −80°C. Following euthanasia, ileum, cecum, and colon contents, as well as spleen and whole brain, were collected, rinsed in cold PBS where appropriate, and frozen at −80°C until analysis. Expanded dissection and specimen-handling details are provided in Supplementary File 4.

### Microbiome profiling

DNA was extracted from ileum, cecum, and colon samples using the Qiagen QIAamp PowerFecal Pro DNA Kit. Bacterial community profiling was performed by amplification of the V4 region of the 16S rRNA gene using primers 515F and 806R, followed by Illumina MiSeq paired-end sequencing (2 × 300 bp). Fungal profiling was performed by ITS2 amplification and MiSeq paired-end sequencing (2 × 300 bp). Reads were quality filtered, denoised, and chimera filtered in QIIME2 using DADA2 to generate amplicon sequence variants (ASVs). Bacterial taxonomy was assigned using a Naive Bayes classifier trained on Greengenes 13_8 at 99% OTUs, and fungal taxonomy was assigned using UNITE-2019.

A subset of 71 gastrointestinal samples was selected for shotgun metagenomic sequencing based on maximal separation in 16S-based unweighted Bray-Curtis ordination space. Libraries were prepared using Nextera XT and sequenced on the Illumina NovaSeq 6000 platform (2 × 150 bp). After quality control and host read removal with KneadData, taxonomic profiling was performed with Kraken2 using the standard NCBI database (confidence threshold 0.3). Functional profiling of genes and pathways was performed with HUMAnN 3.0 using the ChocoPhlAn and UniRef90 databases, with outputs normalized to copies per million. Gut-brain modules (GBMs) were derived by mapping UniRef90 gene families to KEGG orthologs and then to the GBM database using the Omixer-RPM workflow (17). Read depth summaries, processing parameters, and additional bioinformatic details are provided in Supplementary File 4.

### Metabolomics

For fecal metabolomics, pooled week 5–7 stool samples were analyzed for amino acids, lactic acid, short-chain fatty acids, and branched-chain fatty acids by targeted LC-MS. For brain metabolomics, whole-brain tissue from a subset of mice included in the metagenomic analysis was analyzed by targeted LC-MS/MS for selected neurotransmission-related metabolites, including acetylcholine, dopamine, GABA, isoleucine, leucine, nicotinamide, phenylalanine, tryptophan, tyrosine, kynurenine, kynurenic acid, picolinic acid, xanthurenic acid, and serotonin.

Fecal and brain metabolomics data were normalized by sample sum, log transformed, and autoscaled in MetaboAnalyst 5.0. PCA and heatmaps were generated in MetaboAnalyst, and remaining visualizations were produced in R. Extraction, derivatization, instrumentation, and data-processing details are provided in Supplementary File 4.

### Cytokine analysis

Plasma cytokines were quantified at the University of Minnesota Cytokine Reference Laboratory using a multiplex bead-based immunoassay. Samples were analyzed in duplicate using the Luminex Mouse Discovery 8-plex magnetic bead assay (R&D Systems) according to the manufacturer’s protocol and read on an Intelliflex instrument. Concentrations were interpolated from fitted standard curves using Belysa software. Additional assay details are provided in Supplementary File 4.

### Statistical analysis

All analyses were performed in R (v4.5.1) unless otherwise stated. To address reviewer concerns regarding analytical transparency, statistical testing was structured to distinguish primary from exploratory findings.

For behavioral outcomes, normality was assessed using the Shapiro–Wilk test. Student’s t-tests were used only when assumptions were satisfied; otherwise, nonparametric tests were used. Two-group univariate comparisons for alpha diversity, cytokines, relative abundance of specific taxa, metabolite abundances, and other univariate outcomes were assessed using Mann–Whitney U tests. Four-group comparisons were assessed using Kruskal–Wallis tests followed by Dunn tests where appropriate.

Beta diversity was assessed by PCoA and PERMANOVA using unweighted Bray-Curtis (Sørensen-equivalent presence/absence-transformed) and weighted Bray-Curtis distances. Both overall models and within-diet comparisons were examined to evaluate diet, kombucha exposure, and diet × kombucha effects.

Differential abundance testing for taxa, pathways, and genes was performed using MaAsLin3 with total sum scaling normalization. Features with q < 0.1 were treated as FDR-supported primary findings, whereas nominal associations with P < 0.05 but q > 0.1 were interpreted as exploratory. Indicator species analysis was used for fungal features and GBMs when MaAsLin3 did not identify significant candidates.

For metabolomics, data were normalized by sum, log transformed, and autoscaled prior to PCA and heatmap generation. Differential metabolites were tested using Mann-Whitney U or Kruskal–Wallis tests as appropriate.

Exploratory multi-omic integration was performed using CCREPE to identify significant correlations among metagenomic taxa, genes, pathways, GBMs, and fecal metabolites. Chord plots were restricted to positive Spearman correlations > 0.6 among biomarkers associated with kombucha exposure. Because these analyses were exploratory and used relatively permissive q-value thresholds, they were interpreted as hypothesis-generating. Network analyses were visualized in Cytoscape 3.10.3, and network traits included neighborhood connectivity, degree, and average shortest path length. **Expanded software, parameter, and workflow details are provided in Supplementary File 4.**

Significance within groups is indicated as P < 0.05, P < 0.01, and P < 0.001, and trends are indicated by + for P = 0.05-0.09. For analyses involving multiple testing, FDR-adjusted significance was defined as q < 0.1, whereas nominal findings with P < 0.05 but q > 0.1 were considered exploratory.

## Acknowledgements

We would like to thank the University of Minnesota Genomics Center (UMGC) and its pilot grant for supporting sequencing. We thank personnel in the Chen Laboratory, and brain and the Center for Metabolomics and Proteomics (CMSP) at the University of Minnesota for supporting LC-MS/MS analyses. We also acknowledge the Cytokine Reference Laboratory (Dr. Michael Ehrhardt) at the University of Minnesota for their support of serum cytokine analysis. We greatly thank the Minnesota Supercomputing Institute (MSI) for providing a platform to conduct bioinformatics analysis. Finally, we thank Dr. Milena Saqui-Salces (University of Minnesota) for providing useful insights to plan and carry out this work.

## Author Contributions

S.Davison, G. Blanco and K.Guse performed the in vivo mice study. S. Davidson, K. Guse, J.P. Damián and N.Huma performed behavioral analyses. A. Gomez, C. Faulk, S. Davison, K. Guse, A. Sackett, N. Huma and C. Walls performed dissections and DNA extractions. N. Huma and S. Rutschke performed bioinformatic and statistical analysis and visualization. N. Human and A. Gomez wrote the manuscript. C. Chen, J. Zhang sponsored and performed all fecal metabolomic analyses. Conceptualization was done by A.Gomez, C.Faulk and J.P.Damián. All authors helped with editing the manuscript. Supervision and final editing was done by A.Gomez. All authors read and approved the final manuscript.

## Data Availability

The sequence data have been submitted to the Genome Sequence Archive (GSA) databases under accession ID CRA037362 which can be accessed at https://ngdc.cncb.ac.cn/gsa/browse/CRA037362. The online version contains supplementary material which includes additional figures, detailed methods, tables and raw metabolomics data.

## Declarations

### Ethics Statement

The trial was approved by the Institutional Animal Care and Use Committee (IACUC) of University of Minnesota, Protocol ID 2105-39093A. Supplementary materials (figures, methods, tables and metabolomics raw files) may be found in the online DOI.

