## Supplemental Figures for "Diet-dependent effects of kombucha on the gut microbiome and its neuroactive potential: Associations with reduced anxiety and depressive-like behaviors in mice"

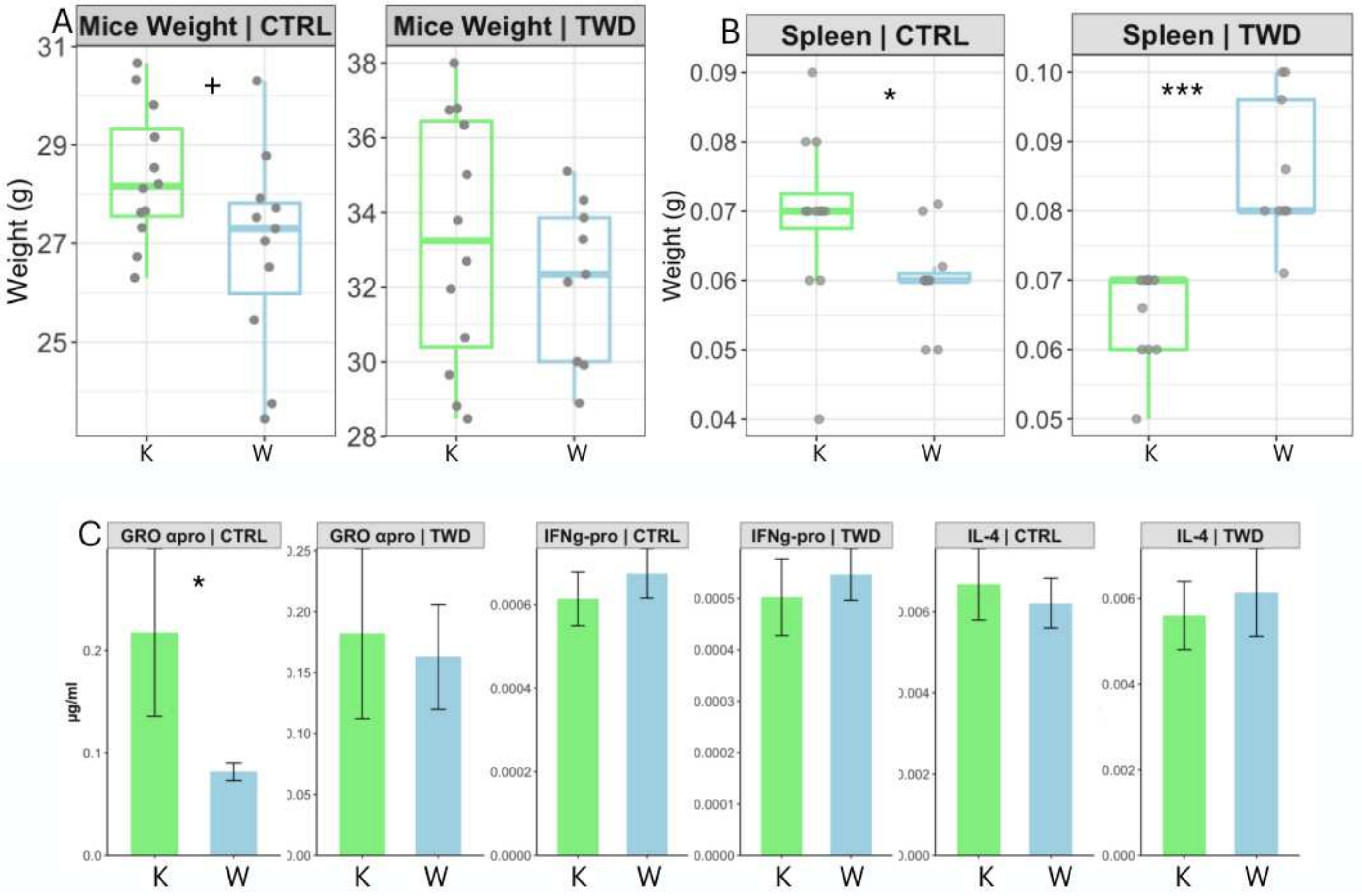


Fig S1. Effects of kombucha on physiological and inflammatory parameters in mice. (A) Body weight. (B) Spleen weight. (C) Serum inflammatory markers. Data are shown as mean ± SEM. Significance for within-diet comparisons was assessed using the Mann–Whitney U test. *P < 0.05, **P < 0.01, and ***P < 0.001; + indicates a trend (P = 0.05-0.09).

### Unweighted Bray Curtis PCoA plot

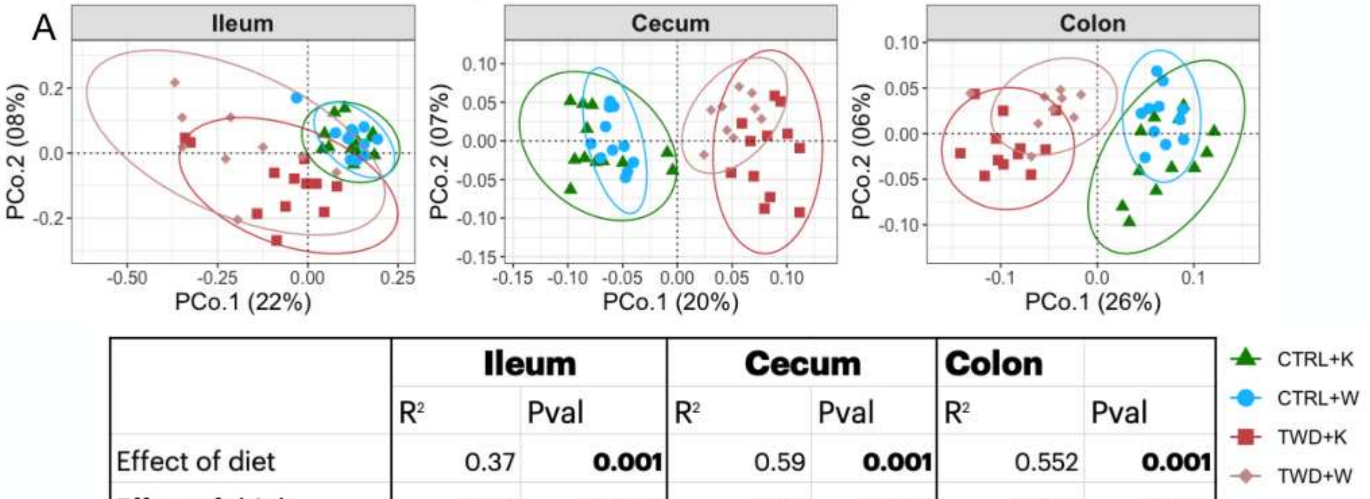


| | Ileum | | Cecum | | Colon | |
|---|---|---|---|---|---|---|
| | $R^2$ | Pval | $R^2$ | Pval | $R^2$ | Pval |
| Effect of diet | 0.37 | **0.001** | 0.59 | **0.001** | 0.552 | **0.001** |
| Effect of drink | 0.06 | **0.005** | 0.02 | 0.073 | 0.014 | 0.222 |
| Effect of interaction | 0.02 | 0.07 | 0.03 | **0.029** | 0.049 | 0.074 |

### Weighted Bray Curtis PCoA plot

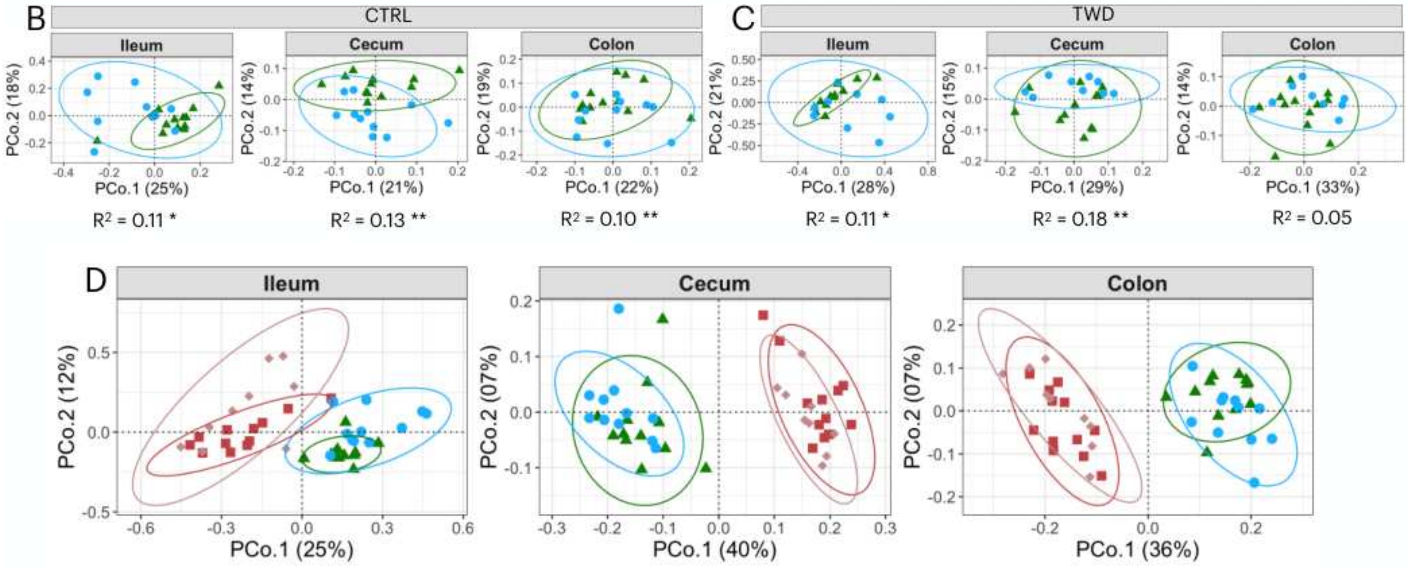


| | Ileum | | Cecum | | Colon | |
|---|---|---|---|---|---|---|
| | $R^2$ | Pval | $R^2$ | Pval | $R^2$ | Pval |
| Effect of diet | 0.37 | **0.001** | 0.59 | **0.001** | 0.552 | **0.001** |
| Effect of drink | 0.06 | **0.007** | 0.02 | 0.087 | 0.014 | 0.207 |
| Effect of interaction | 0.02 | 0.072 | 0.03 | **0.03** | 0.023 | 0.071 |

Fig S2. Diet-dependent effects of kombucha on the gut microbiome. (A) Beta diversity of bacterial communities assessed by PCoA based on unweighted Bray-Curtis (Sørensen-equivalent presence/absence-transformed) distances with PERMANOVA. (B-D) Beta diversity based on weighted Bray-Curtis distances, including overall and within-diet comparisons. Ellipses represent 95% confidence intervals for each group.

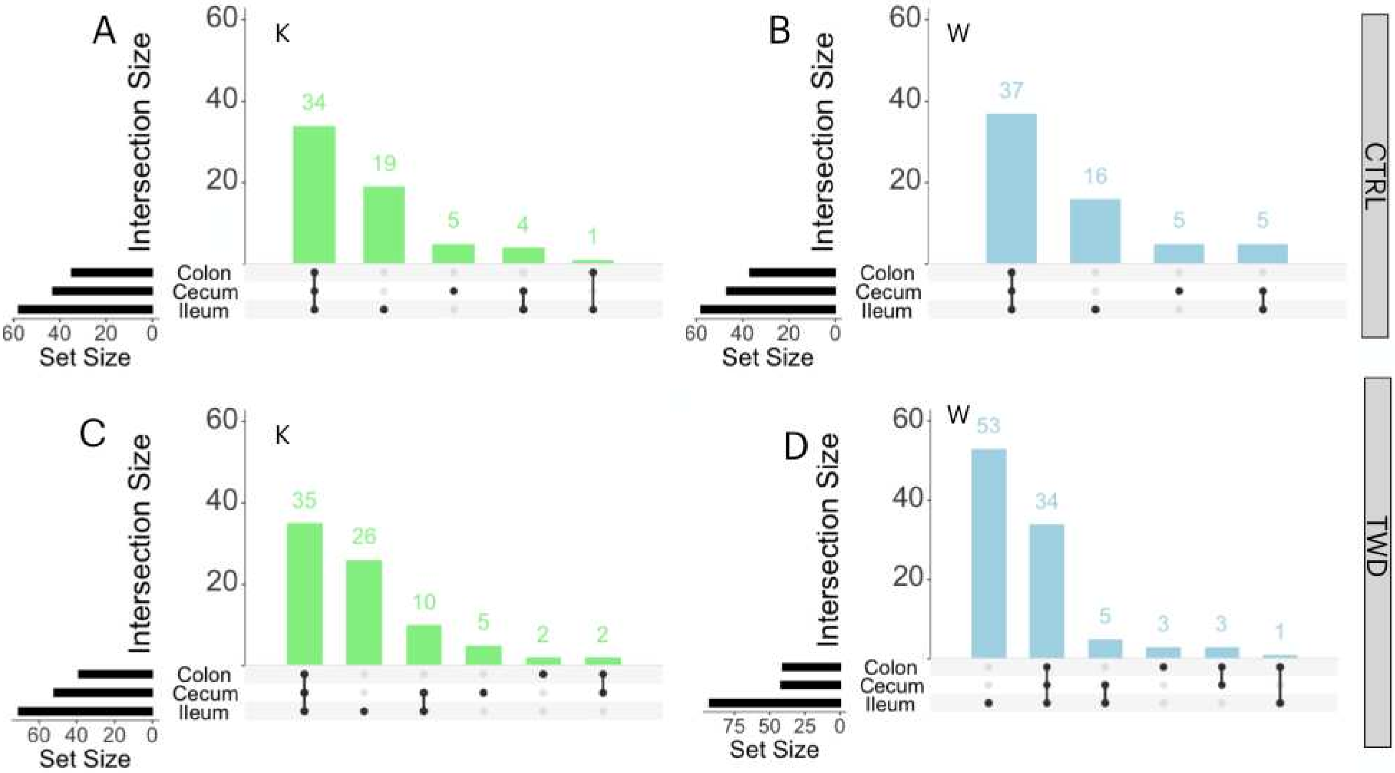


Fig S3. Diet-dependent modulation of bacterial ASV overlap by kombucha. UpSet plots showing intersections of bacterial amplicon sequence variants (ASVs) among ileum, cecum, and colon under CTRL and TWD in the 16S rRNA dataset. Set size indicates the total number of ASVs detected at each gastrointestinal site, and intersection size indicates the number of ASVs shared across sites.

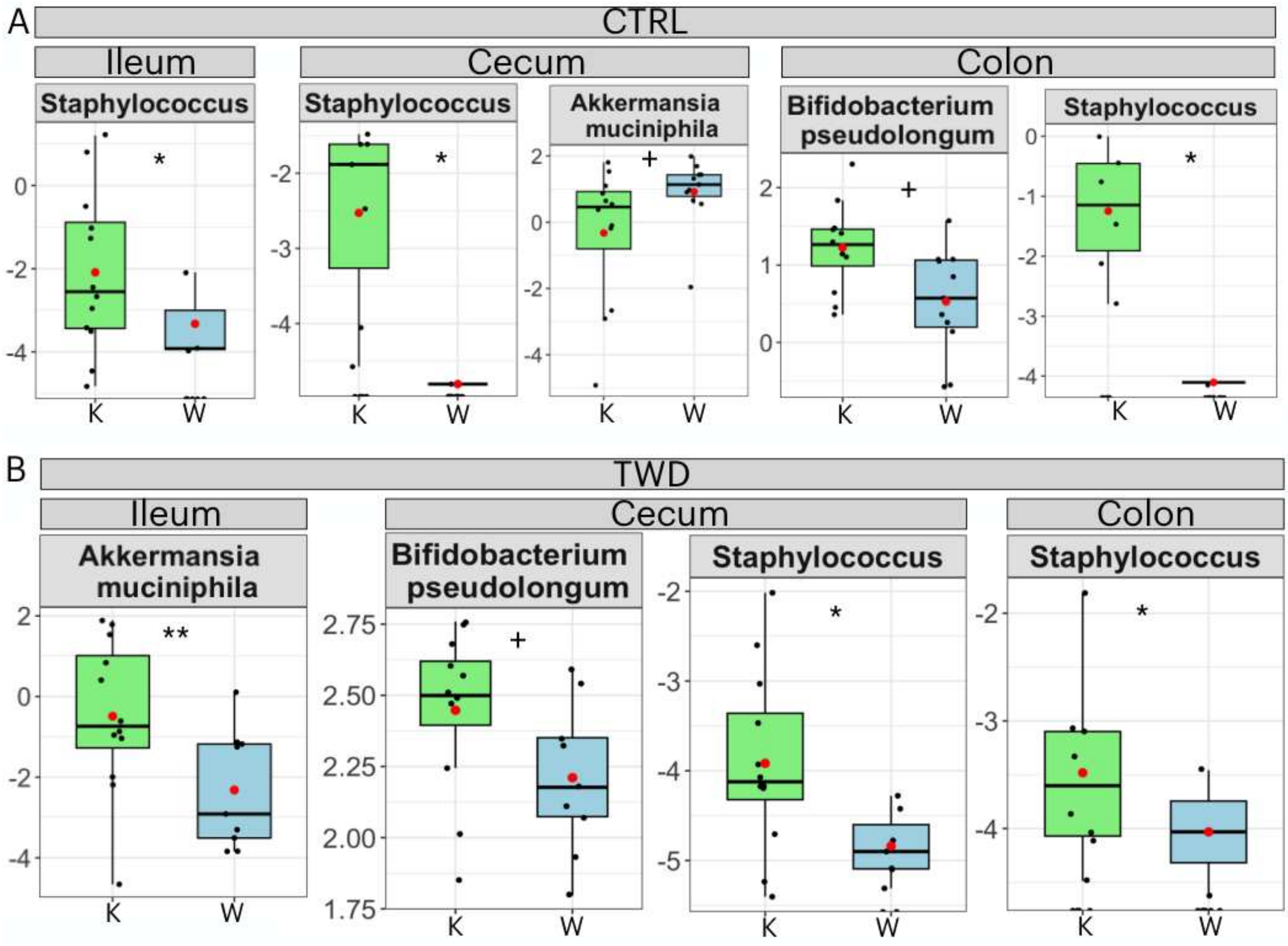


Fig S4. Effects of kombucha on gastrointestinal mycobiome alpha diversity. (A-B) Chao1 richness and Shannon diversity of fungal communities in the ileum, cecum, and colon. Significance for within-diet comparisons was assessed using the Mann–Whitney U test. *P < 0.05, **P < 0.01, and ***P < 0.001.

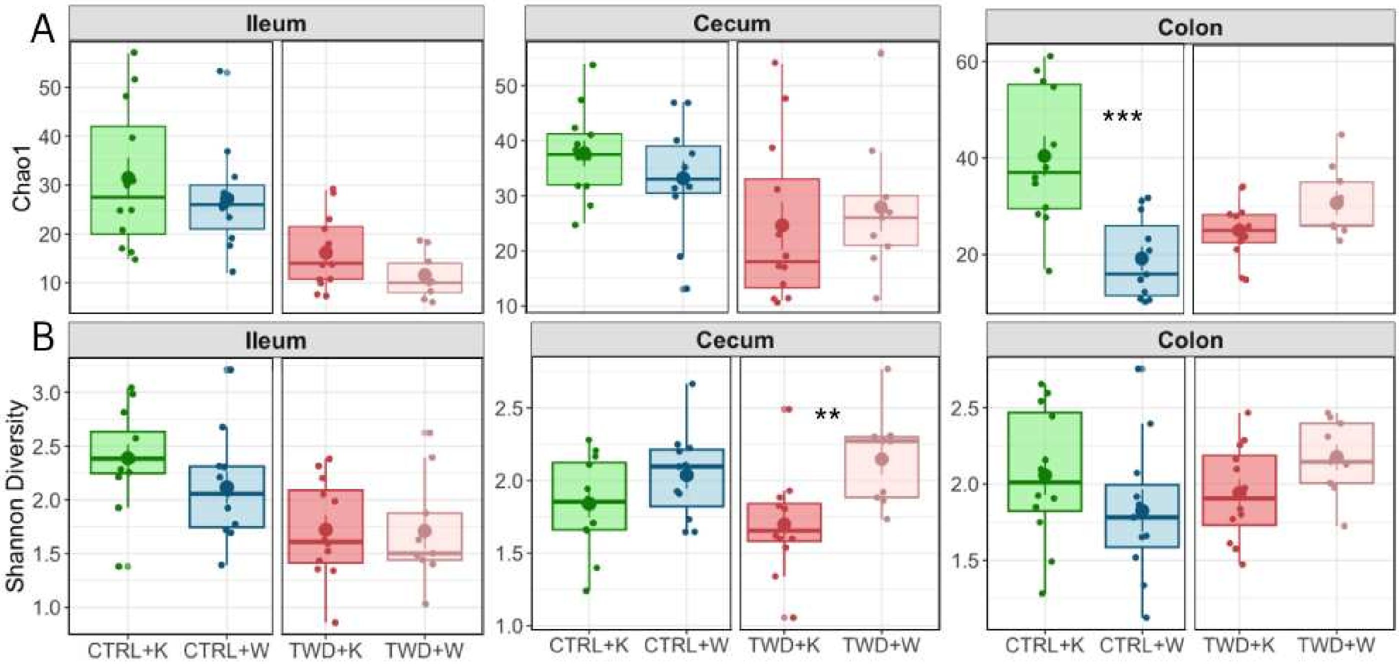


Fig S5. Diet-dependent effects of kombucha on gut mycobiome beta diversity. (A-B) Beta diversity of fungal communities assessed by PCoA based on unweighted and weighted Bray-Curtis distances, respectively, with PERMANOVA. (C-D) Within-diet fungal community comparisons assessed by PCoA based on unweighted and weighted Bray-Curtis distances, respectively. Ellipses represent 95% confidence intervals for each group.

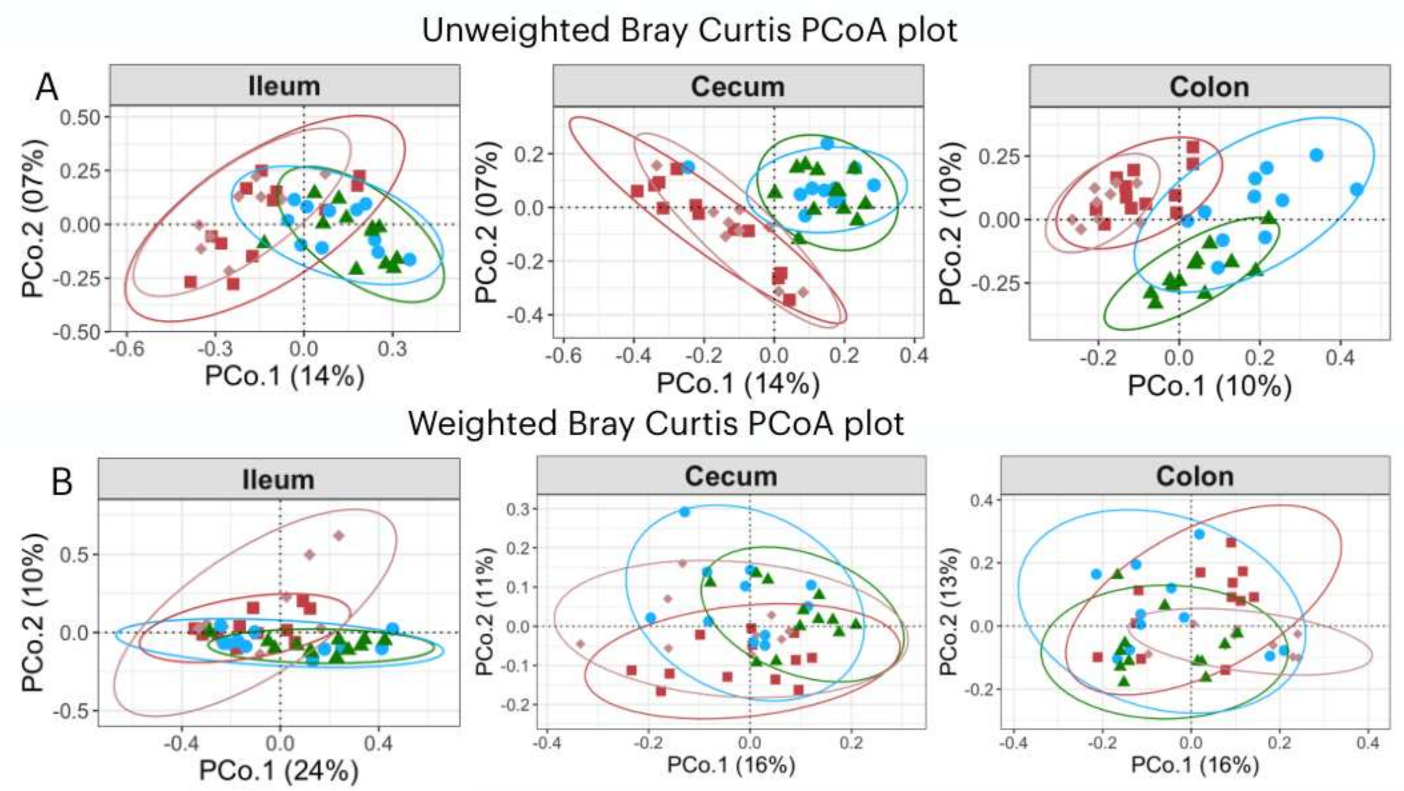


| | Unweighted | | | | | | Weighted | | | | | |
|---|---|---|---|---|---|---|---|---|---|---|---|---|
| | Ileum | | Cecum | | Colon | | Ileum | | Cecum | | Colon | |
| | $R^2$ | Pval | $R^2$ | Pval | $R^2$ | Pval | $R^2$ | Pval | $R^2$ | Pval | $R^2$ | Pval |
| Effect of diet | 0.1 | **0.001** | 0.12 | **0.001** | 0.122 | **0.001** | 0.09 | **0.001** | 0.1 | **0.001** | 0.09 | **0.001** |
| Effect of drink | 0.01 | 0.825 | 0.02 | 0.248 | 0.03 | **0.039** | 0.02 | 0.263 | 0.04 | **0.005** | 0.02 | 0.286 |
| Effect of interaction | 0.01 | 0.636 | 0.03 | 0.038 | 0.05 | **0.001** | 0.02 | 0.407 | 0.04 | **0.016** | 0.06 | **0.001** |

CTRL+K
CTRL+W
TWD+K
TWD+W

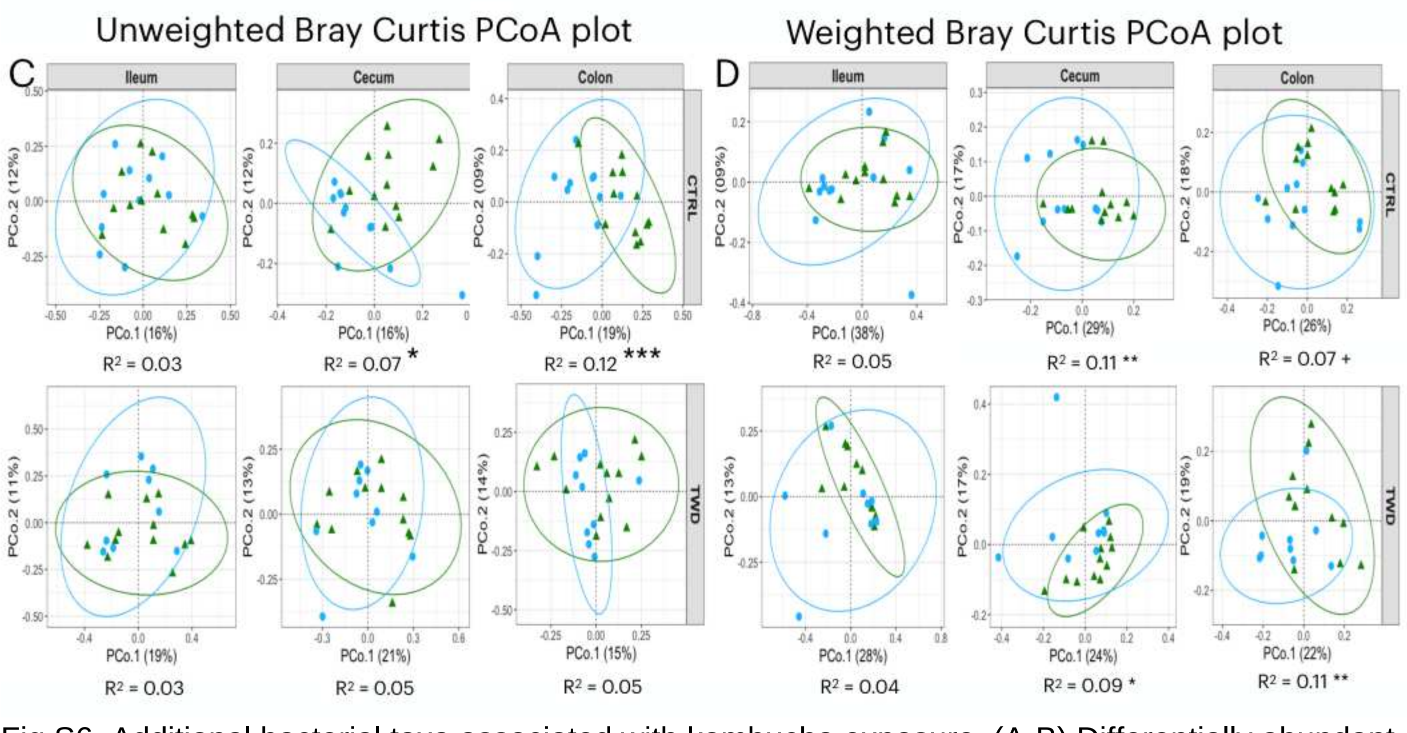


Fig S6. Additional bacterial taxa associated with kombucha exposure. (A-B) Differentially abundant bacterial features identified by MaAsLin3 in within-diet comparisons. These signals were nominal rather than FDR-supported ($P < 0.05$, q = 0.16-0.4). *$P < 0.05$, **$P < 0.01$, and ***$P < 0.001$; + indicates a trend ($P$ = 0.05-0.09).

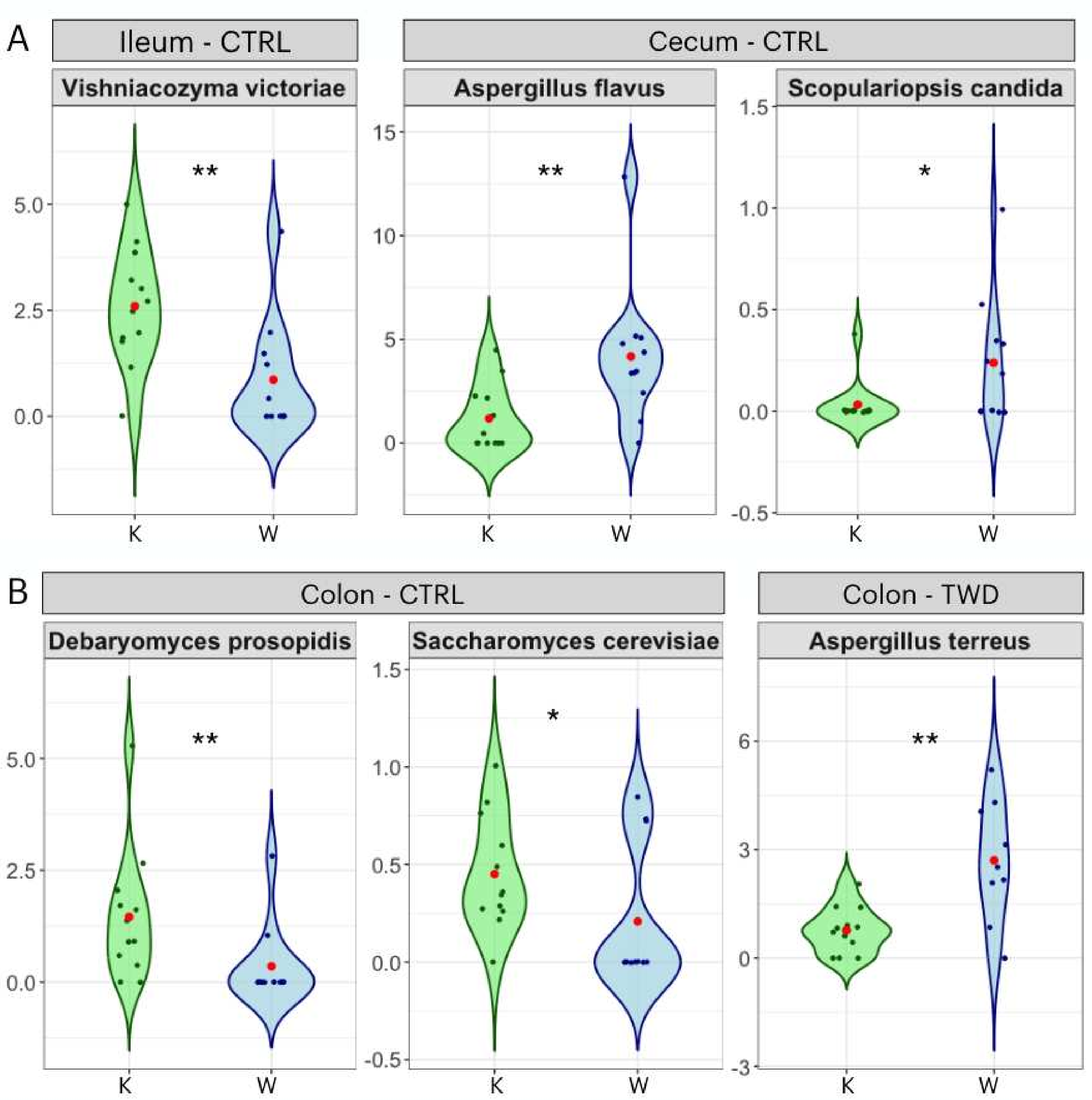


Fig S7. Diet- and site-specific fungal taxa associated with kombucha exposure. (A-B) Differentially abundant fungal features from ITS2 data identified by indicator species analysis (IndVal; $P < 0.05$, indicator value > 0.6). *$P < 0.05$, **$P < 0.01$, and ***$P < 0.001$.

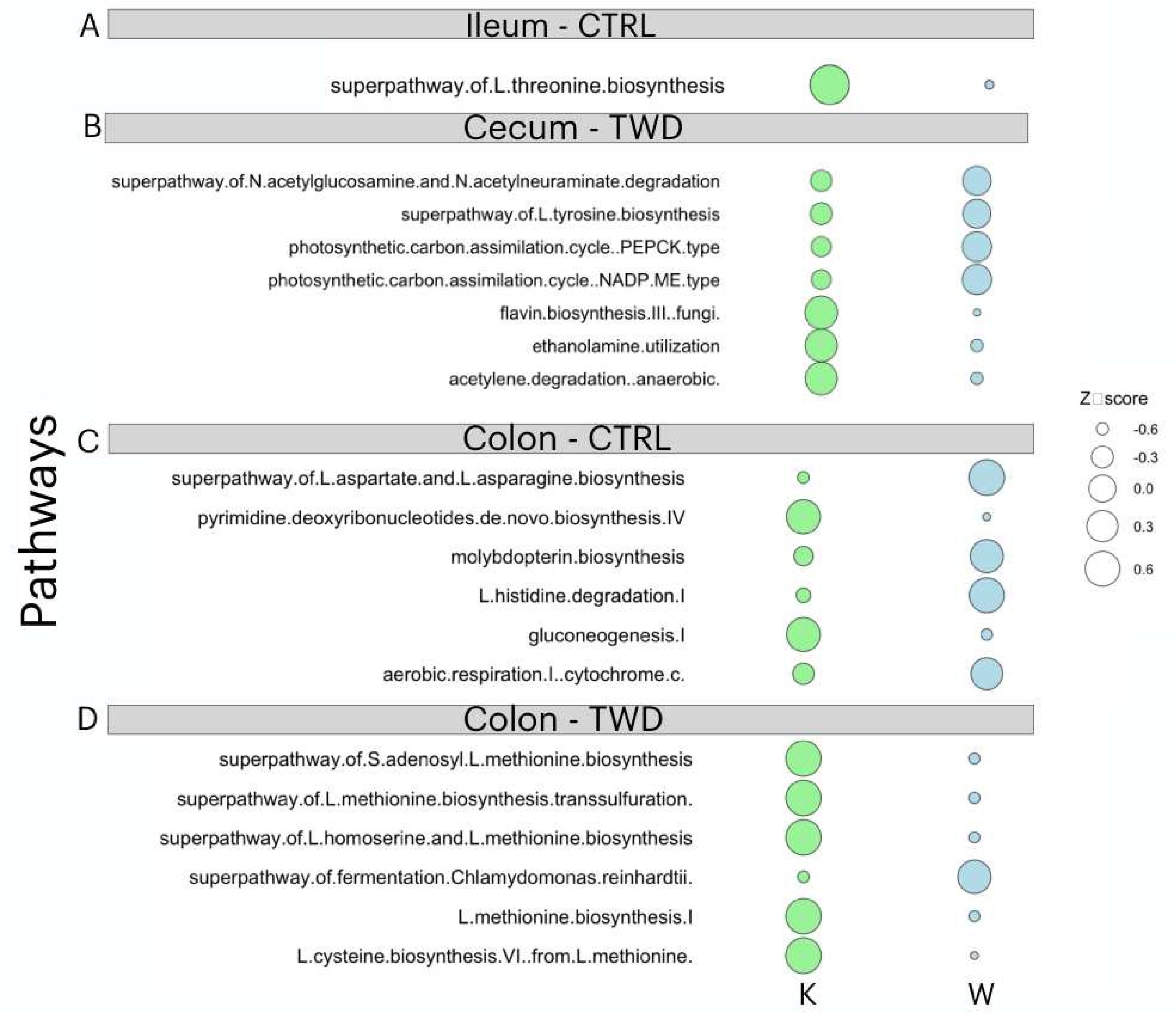


Fig S8. Additional pathway shifts in the gut microbiome associated with kombucha exposure. (A-D) Differentially abundant microbial pathways from shotgun metagenomic data identified by MaAsLin3. These pathway-level signals were nominal rather than FDR-supported (P < 0.05, q = 0.7-0.9).

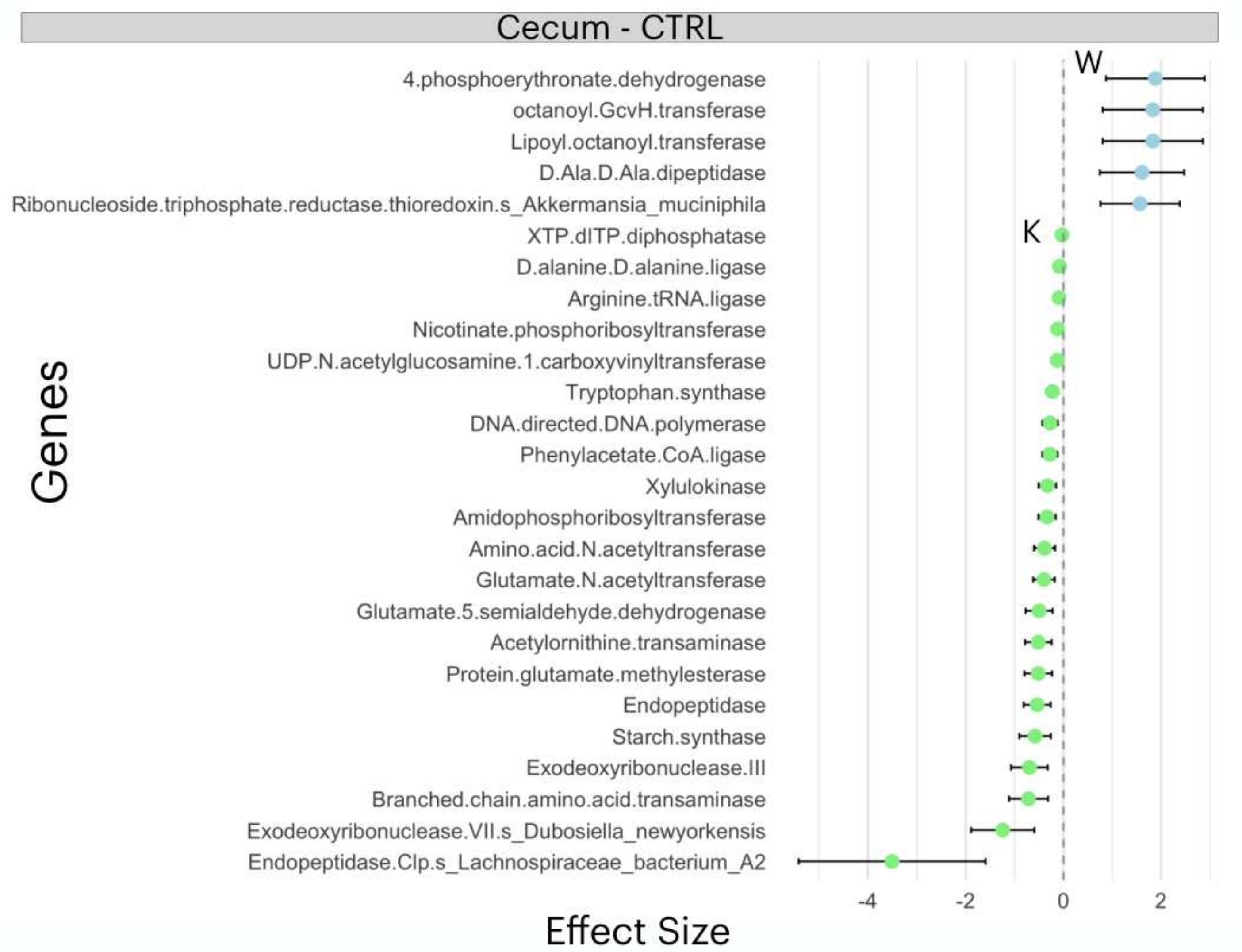


Fig S9. Additional microbial genes associated with kombucha exposure. Effect sizes of differentially abundant microbial genes from metagenomic data identified by MaAsLin3 in the cecum under CTRL and other site-specific contrasts (P < 0.05, q < 0.1 unless otherwise indicated).

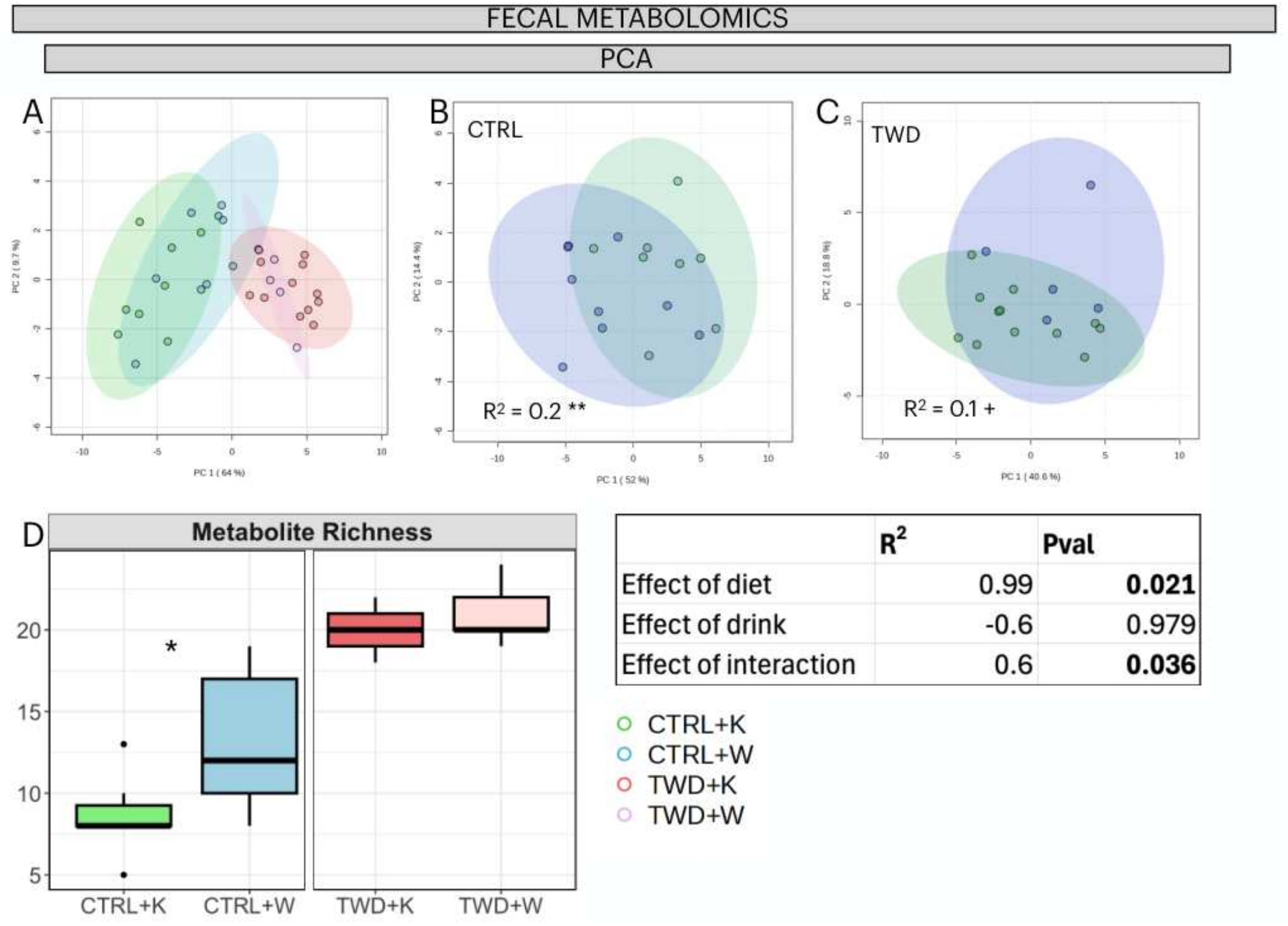


| | $R^2$ | Pval |
|---|---|---|
| Effect of diet | 0.99 | **0.021** |
| Effect of drink | -0.6 | 0.979 |
| Effect of interaction | 0.6 | **0.036** |

Fig S10. Diet-dependent fecal metabolite profiles with kombucha exposure. (A) Multivariate analysis of fecal metabolite structure assessed by PCA/PERMANOVA, showing the effects of diet and drink. (B-C) Within-diet fecal metabolite structure for CTRL and TWD, respectively. Data were normalized by sum, log transformed, and autoscaled. (D) Fecal metabolite richness across diet and drink groups. *P < 0.05, **P < 0.01, and ***P < 0.001 for within-diet comparisons; + indicates a trend (P = 0.05-0.09). Inset tables report $R^2$ and P values for the multivariate models.

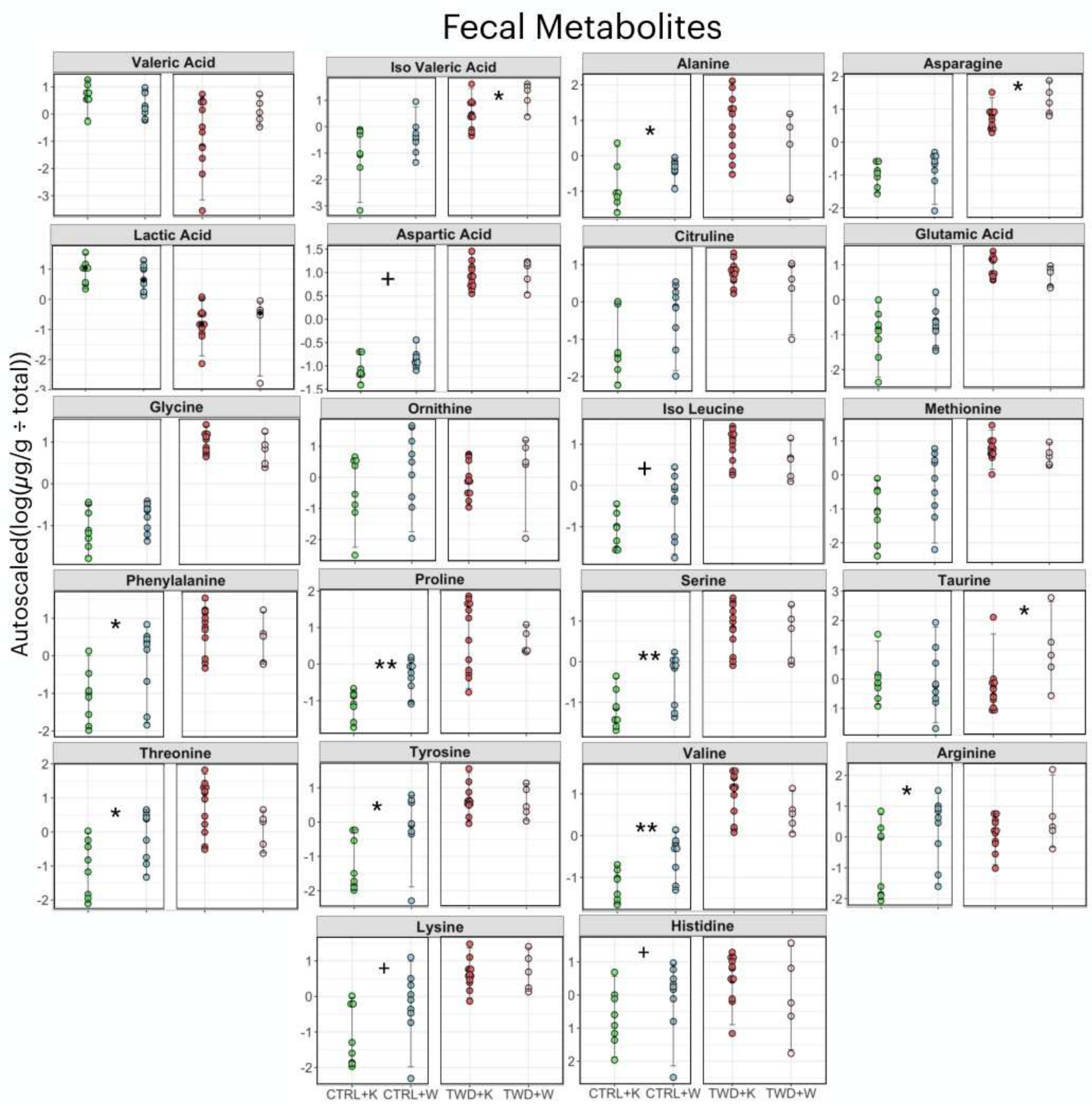


Fig S11. Additional fecal metabolites associated with kombucha exposure across diets. Beeswarm plots showing fatty acids and amino acids not displayed in the main figures. Significance for within-diet comparisons was assessed using the Mann–Whitney U test. *P < 0.05, **P < 0.01, and ***P < 0.001; + indicates a trend (P = 0.05-0.09).

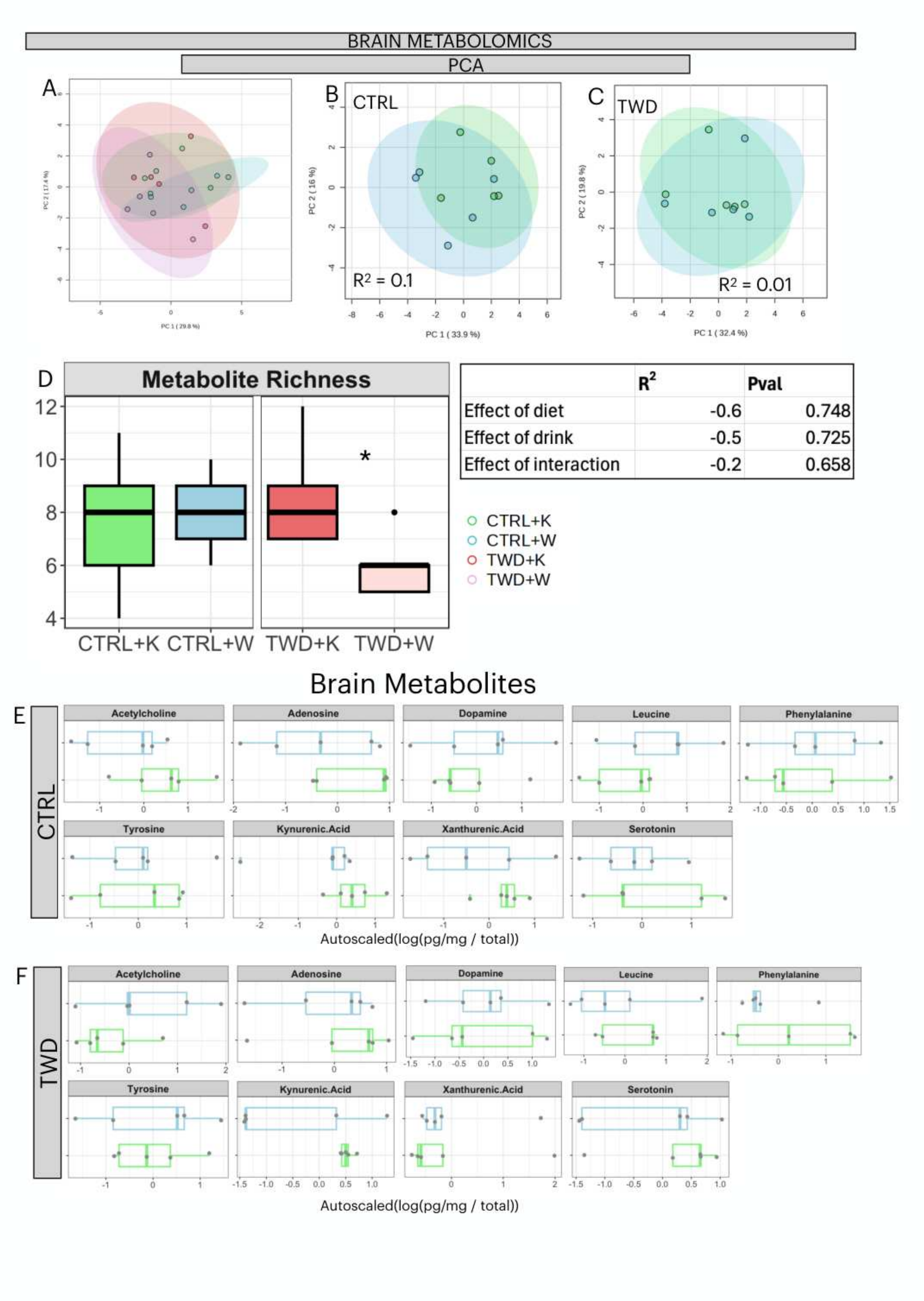


Fig S12. Diet-dependent whole-brain metabolite profiles with kombucha exposure. (A) Multivariate analysis of whole-brain metabolite structure assessed by PCA/PERMANOVA, showing the effects of diet and drink. (B-C) Within-diet whole-brain metabolite structure for CTRL and TWD, respectively. Data were normalized by sum, log transformed, and autoscaled. (D) Whole-brain metabolite richness across diet and drink groups. No significant effects were detected. Inset tables report $R^2$ and P values for the multivariate models. (E-F) Additional whole-brain metabolites that did not differ significantly between groups ($P > 0.05$).

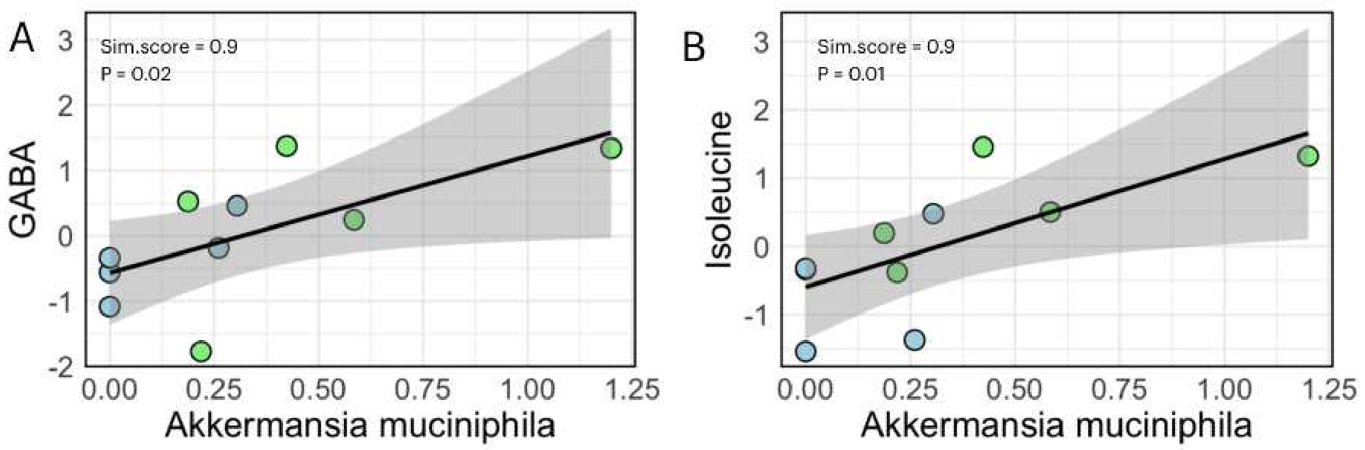


Fig S13. Associations between ileal Akkermansia muciniphila and brain metabolites under TWD. (A-B) Scatter plots showing positive correlations between ileal Akkermansia muciniphila abundance and whole-brain GABA (A) or isoleucine (B) under TWD.

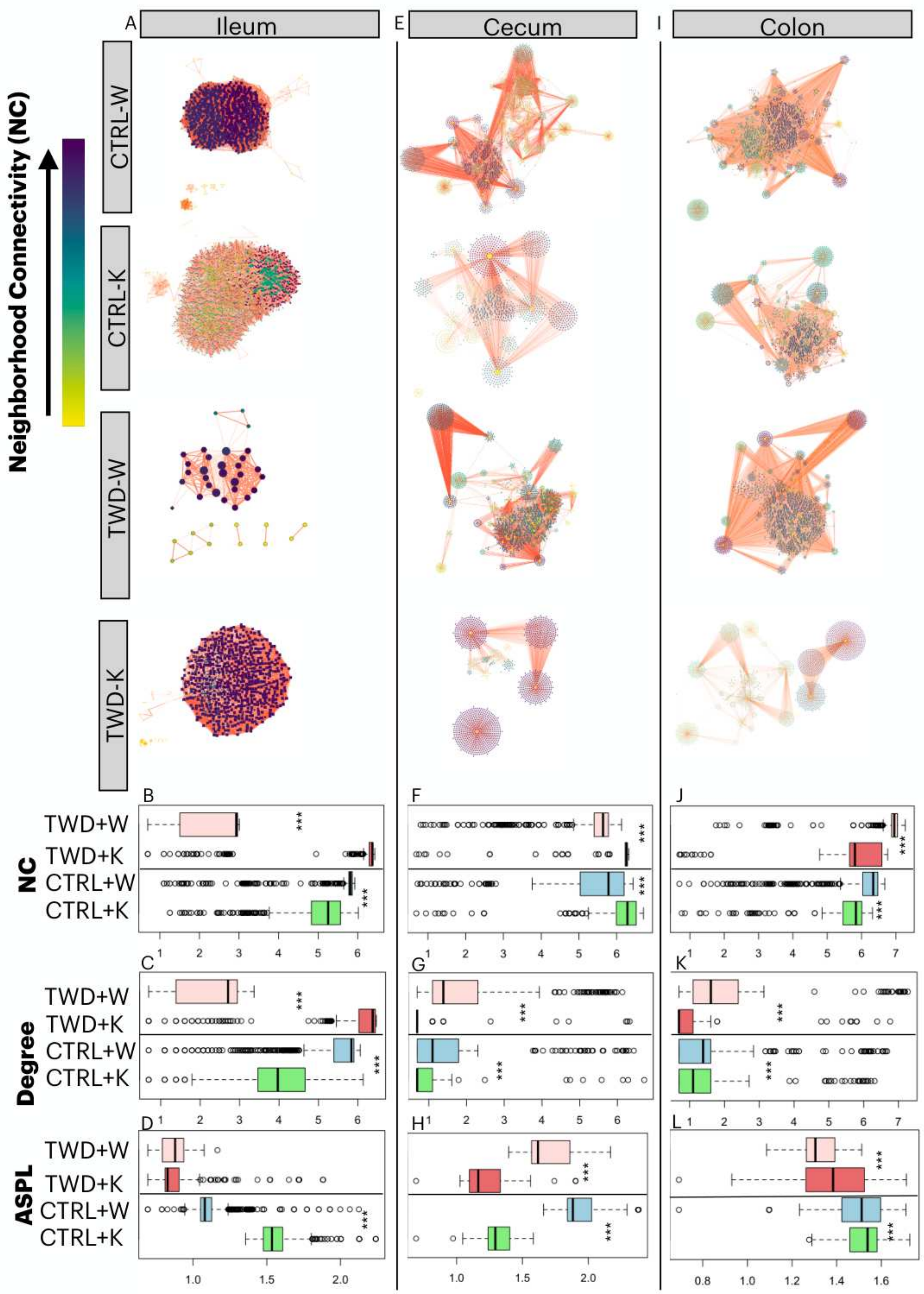


Fig S14. Co-abundance networks linking metagenomic features across gut sites. (A-D) Ileal co-abundance network analyses linking taxonomy, genes, and gut-brain modules (GBMs), including network plots and comparisons of neighborhood connectivity (B), degree (C), and average shortest path length (ASPL) (D) across diet and drink groups. (E-H) Equivalent analyses for the cecum. (I-L) Equivalent analyses for the colon. Node color represents neighborhood connectivity and edge color represents correlation strength. *P < 0.05, **P < 0.01, and ***P < 0.001 denote significant within-diet differences between kombucha and water.

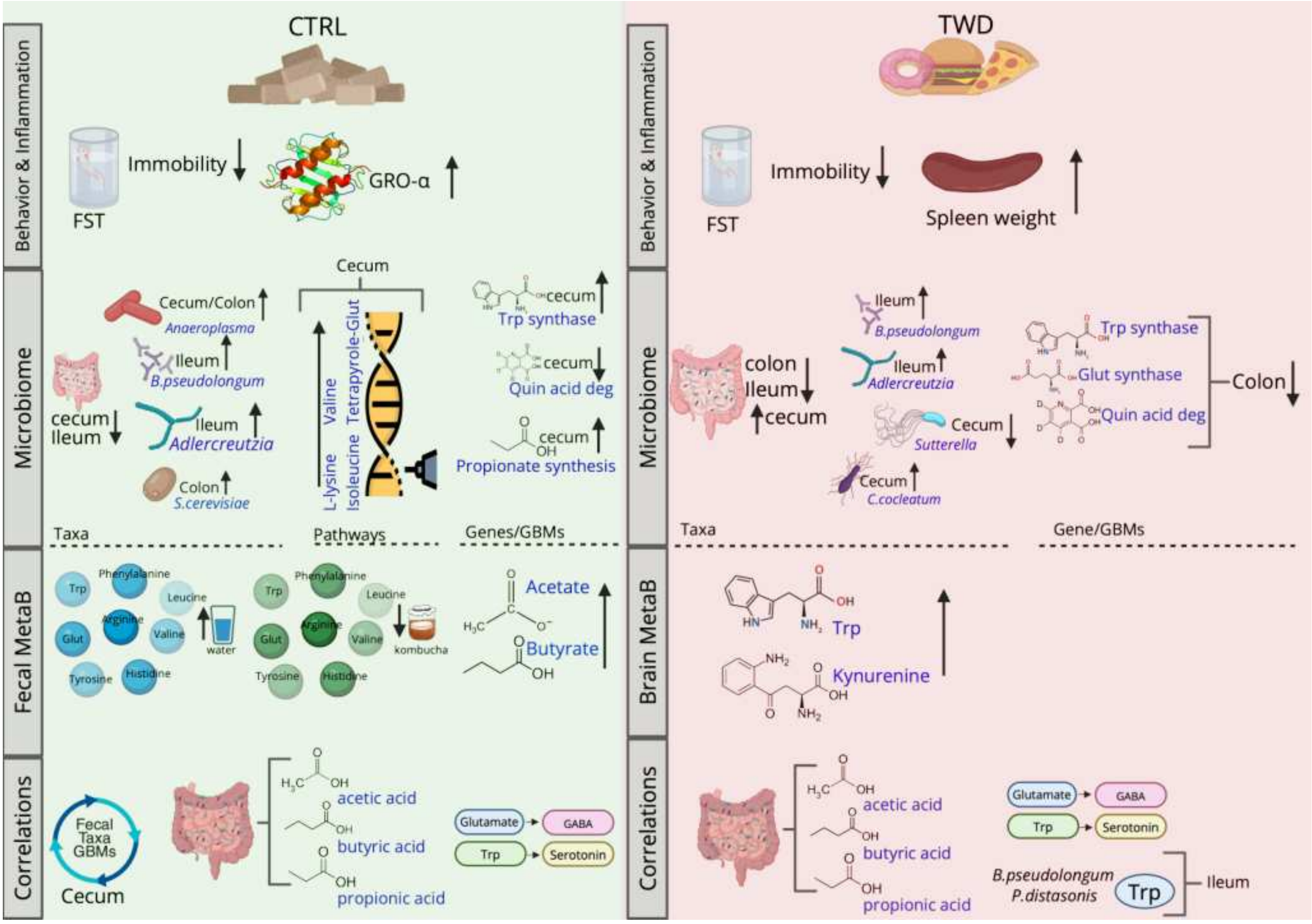


Fig S15. Schematic overview of study findings. Illustration summarizing the main results of the study, highlighting diet-dependent effects of kombucha on gut microbial composition, microbial functional potential, fecal and brain metabolite profiles, and behavior. Created with BioRender.
