## Supplementary File 4 for "Diet-dependent effects of kombucha on the gut microbiome and its neuroactive potential: Associations with reduced anxiety and depressive-like behaviors in mice"

**SUPPLEMENTARY METHODS**

**S1. Preparation of kombucha tea**

Kombucha was prepared from sencha green tea using a standardized fermentation protocol. Briefly, organic cane sugar was dissolved in 1 L of water heated to 80°C, followed by infusion of 10 g sencha green tea for 10 min. After cooling to room temperature (25°C), the tea was transferred to a sterile 2 L glass jar and inoculated with 100 mL starter kombucha plus a symbiotic culture of bacteria and yeast (SCOBY). Fermentation proceeded aerobically at 25°C for 21 days under a breathable cloth cover. The fermented kombucha was then stored at 4°C until use. Extended fermentation was selected to promote organic acid production, reduce residual sugar, and maintain pH between 2.5 and 3.5.

**S2. Endpoint dissection and specimen handling**

Stool samples were collected weekly from weeks 1–7 and stored at −80°C. For fecal metabolomics, samples from weeks 5–7 were pooled within mouse before analysis. Approximately 50 µL of blood was collected from the saphenous vein at the end of the study into serum separator tubes, centrifuged at 1,200 × g for 10 min, aliquoted, and stored at −80°C.

Following euthanasia, ileum, cecum, and colon contents, as well as spleen and whole brain, were collected following standard rodent dissection procedures (1,2). Briefly, euthanized mice were placed dorsally and the abdominal cavity was exposed by a midline incision. The colon was isolated near the rectum/anus, the cecum was excised at the ileocecal junction, and the ileum was isolated proximal to the cecum. Excess mesenteric and adipose tissue were removed before freezing. The spleen was excised from the posterior side of the stomach beneath the rib cage. Whole brains were harvested by opening the skull with micro-scissors and carefully separating the intact brain tissue from the cranial cavity (2). Digestive tissues and brains were rinsed in cold PBS where appropriate, transferred to pre-labeled 1.5 mL tubes, and stored at −80°C until downstream analysis.

**S3. 16S rRNA and ITS2 amplicon sequencing and processing**

Bacterial community composition was profiled by amplifying the V4 hypervariable region of the 16S rRNA gene using primers 515F (GTGYCAGCMGCCGCGGTAA) and 806R (GGACTACNVGGGTWTCTAAT) with a dual-index library preparation approach. Fungal community composition was profiled by amplifying ITS2 using primers GAACGCAGCGAAATGCGA and GTGAATCATCGAATCTTTG. Pooled libraries were sequenced at the University of Minnesota Genomics Center on the Illumina MiSeq platform using paired-end 2 × 300 bp chemistry.

Raw reads were processed to remove primers and low-quality bases (PHRED < 30), then denoised and chimera filtered using the DADA2 plugin within QIIME2 (3,4). For 16S data, the average raw sequencing depth was 57,705 reads/sample (range 23,935–106,619), and the average post-filter depth was 41,964 reads/sample (range 14,664–84,156), corresponding to 72.7% retention. Taxonomy was assigned using a pretrained Naive Bayes classifier trained on the Greengenes 13_8 database at 99% OTUs.

For ITS2 data, the average raw sequencing depth was 37,201 reads/sample (range 3,147–81,263), and the average post-filter depth was 19,329 reads/sample (range 1,341–42,492), corresponding to 53% read retention. Fungal taxonomy was assigned using the UNITE-2019 database. Resulting bacterial and fungal ASV tables were used for downstream ecological and differential abundance analyses.

**S4. Shotgun metagenomic sequencing and processing**

Shotgun metagenomic profiling was performed on 71 gastrointestinal samples from ileum, cecum, and colon. This subset was selected based on maximal separation of samples in 16S-based unweighted Bray-Curtis ordination space. Libraries were prepared using the Nextera XT DNA library preparation kit (Illumina) and sequenced on the Illumina NovaSeq 6000 platform using paired-end 2 × 150 bp reads.

The average number of reads remaining after initial quality control was 2,162,409 reads/sample (range 1,051,423–18,714,292). Reads were processed with KneadData using SLIDINGWINDOW:4:40 and MINLEN:90. After host read removal, the average number of reads remaining was 587,756/sample (range 142,071–4,762,763). Taxonomic assignment was performed with Kraken2 using the standard NCBI database and a confidence threshold of 0.3. Functional profiling of pathways and genes was performed with HUMAnN 3.0 using the ChocoPhlAn nucleotide and UniRef90 protein databases. Gene and pathway abundances were normalized to copies per million.

For gut-brain module (GBM) analysis, UniRef90 gene families detected by HUMAnN 3.0 were regrouped to KEGG orthologs and mapped to the GBM database using the Omixer-RPM workflow as previously described (5). GBMs represent KEGG ortholog groups associated with the synthesis or degradation of neuroactive compounds.

**S5. Metabolomics analysis**

**S5.1. Sample collection and targeted panels**

Fecal samples collected from weeks 5–7 were pooled per mouse and stored at −80°C until analysis. Approximately 50 mg stool was transferred to 1.5 mL tubes and shipped on dry ice to the Chen Laboratory (University of Minnesota) for targeted LC-MS analysis of amino acids, lactic acid, short-chain fatty acids, and branched-chain fatty acids.

Whole-brain tissue from a subset of mice included in the metagenomic analysis was analyzed at the Center for Metabolomics and Proteomics (University of Minnesota) for targeted LC-MS/MS quantification of selected neurotransmission-related metabolites, including acetylcholine, dopamine, GABA, isoleucine, leucine, nicotinamide, phenylalanine, tryptophan, tyrosine, kynurenine, kynurenic acid, picolinic acid, xanthurenic acid, and serotonin.

**S5.2. Fecal metabolite extraction and LC-MS**

Fecal samples were extracted in 50% aqueous acetonitrile (1:10, w/v), centrifuged at 18,000 × g for 10 min, and supernatants were collected. SCFAs were derivatized with 2-hydrazinoquinoline and analyzed in positive ion mode by LC-MS. Metabolites containing amino or hydroxyl groups were derivatized with dansyl chloride by mixing 5 µL sample or standard, 5 µL 100 µM d5-tryptophan internal standard, 50 µL 10 mM sodium carbonate, and 100 µL dansyl chloride solution (3 mg/mL in acetone), followed by incubation at 60°C for 15 min, centrifugation, and analysis of the supernatant.

LC-MS was performed on a Waters UPLC-QTOF-MS system with a BEH C18 column using a 10 min water–acetonitrile gradient containing 0.1% formic acid (6–9). Electrospray ionization was carried out in positive mode with a 3 kV capillary voltage and 30 V cone voltage. Source and desolvation temperatures were 120°C and 350°C, respectively. Nitrogen was used as cone and desolvation gas, and argon as the collision gas. Calibration was performed with sodium formate and leucine-enkephalin. Quantification was based on analyte/internal standard ratios relative to external standard curves in MassLynx and QuanLynx.

**S5.3. Brain metabolite extraction and LC-MS/MS**

Brain tissues were weighed and homogenized in 0.1% ascorbic acid plus 0.5% phosphoric acid (1:2, w/v) for 3 × 30 s at 8,000 rpm and 6°C. Homogenates were centrifuged at 14,000 rpm for 20 min at 4°C, and 200 µL supernatant was mixed with 20 µL isotope-labeled internal standard mixture. For phospholipid extraction, 660 µL of 1% formic acid in acetonitrile (3:1, v/v) was added, samples were incubated at −20°C for 2 h, filtered through Waters Ostro plates, dried under nitrogen, and reconstituted in 100 µL 0.1% formic acid in water.

Samples were analyzed using a Waters Acquity Premier UHPLC coupled to a Sciex QTRAP 6500. Separation was performed on an Acquity Premier BEH C18 column (2.1 × 100 mm, 1.7 µm) at 0.2 mL/min with a 10 µL injection. Mobile phase A was 0.1% formic acid in water and mobile phase B was 0.1% formic acid in acetonitrile. The gradient was 0.1–5 min (98–65% A), 5–11 min (65–2% A), 11–12 min hold (2% A), 12–13.5 min (2–98% A), and 13.5–15 min hold (98% A). Data were imported into Skyline v23.1.0.380 for quantification.

**S5.4. Metabolomics data processing**

Fecal and brain metabolomics data were normalized by sample sum, log transformed, and autoscaled in MetaboAnalyst 5.0 (15). Principal component analysis and heatmaps were generated in MetaboAnalyst, while all other visualizations were produced in R.

**S6. Cytokine analysis**

Plasma cytokines were quantified at the University of Minnesota Cytokine Reference Laboratory using the Luminex Mouse Discovery 8-plex magnetic bead assay (R&D Systems; cat. #LXASHM-08) according to the manufacturer’s protocol (10). Samples were run in duplicate. Briefly, fluorescently coded magnetic beads coated with analyte-specific capture antibodies were incubated with plasma samples, followed by detection with biotinylated antibodies and phycoerythrin-conjugated streptavidin. Beads were read on an Intelliflex dual-laser instrument, with one laser identifying bead class and the other measuring PE signal intensity. Concentrations were interpolated from 4- or 5-parameter fitted standard curves using Belysa Immunoassay Curve-Fitting Software.

**S7. Extended statistical workflow**

Microbiome analyses were conducted in R (v4.5.1), primarily using vegan and ape for ecological analyses (11,12). Most analyses were performed as within-diet comparisons of kombucha versus water in CTRL or TWD, although diet × drink interactions were also evaluated for selected endpoints.

Behavioral outcomes were tested using Student’s t-test when Shapiro–Wilk tests supported approximate normality. Two-group univariate comparisons, including alpha diversity, cytokines, and relative abundance of selected taxa or metabolites, were assessed using Mann–Whitney U tests. Four-group comparisons were assessed using Kruskal–Wallis tests followed by Dunn tests where appropriate.

Beta diversity was evaluated by PCoA and PERMANOVA. Differentially abundant taxa, pathways, and genes were identified using MaAsLin3 with total sum scaling normalization and FDR correction at q < 0.1 (13). Indicator species analysis was implemented with labdsv when appropriate (14). In the revised manuscript, q < 0.1 findings were treated as primary, whereas nominal P < 0.05 findings with q > 0.1 were treated as exploratory.

Metabolomics data were normalized by sum, log transformed, and autoscaled in MetaboAnalyst 5.0 (15). PCA and heatmaps were generated in MetaboAnalyst, and differential metabolites were subsequently tested using Mann–Whitney U or Kruskal–Wallis tests as appropriate.

Exploratory multi-omic associations among taxa, genes, pathways, gut-brain modules, and fecal metabolites were evaluated using CCREPE (16). Targeted association plots were restricted to positive Spearman correlations > 0.6 among biomarkers associated with kombucha exposure. Chord diagrams were generated using ComplexHeatmap and circlize (17,18). Network analyses were visualized in Cytoscape 3.10.3, and summary topology metrics included neighborhood connectivity, degree, and average shortest path length. Because these integrative analyses were exploratory, they were interpreted as hypothesis-generating rather than confirmatory.

Significant within-group differences are denoted in figures as *P < 0.05, **P < 0.01, and ***P < 0.001, while trends are denoted by + for P = 0.05–0.09.
