## Supplementary File 5 for "Diet-dependent effects of kombucha on the gut microbiome and its neuroactive potential: Associations with reduced anxiety and depressive-like behaviors in mice"

**ABBREVIATIONS**

| BBB | blood brain barrier |
| --- | --- |
| FST | forced swimming test |
| MBT | marble burying test |
| GRO-α | growth regulating oncogene |
| CTRL | Control diet (chow) |
| TWD | Total western diet |
| GBMs | Gut brain modules |
| Trp | Tryptophan |
| GABA | Y-gamma aminobutyric acid |
| SCOBY | Symbiotic culture of bacteria and yeast |
| SCFAs | Short chain fatty acids |
| GI | Gastrointestinal |
| LS-MS | liquid chromatography-mass spectrometry |
| ELISA | Enzyme linked immunosorbent assay |
| AA | Amino acids |
| ASV | Amplicon Sequence Variants |
| FFAR3 | Free fatty acid receptor 3 |
| Kyn | Kynurenine |
| QUIN | Quinolinic |
| IV | Indicator value |
